# SPACe (Swift Phenotypic Analysis of Cells): an open-source, single cell analysis of Cell Painting data

**DOI:** 10.1101/2024.03.21.586132

**Authors:** Fabio Stossi, Pankaj K. Singh, Michela Marini, Kazem Safari, Adam T. Szafran, Alejandra Rivera Tostado, Christopher D. Candler, Maureen G. Mancini, Elina A. Mosa, Michael J. Bolt, Demetrio Labate, Michael A. Mancini

## Abstract

Phenotypic profiling by high throughput microscopy has become one of the leading tools for screening large sets of perturbations in cellular models. Of the numerous methods used over the years, the flexible and economical Cell Painting (CP) assay has been central in the field, allowing for large screening campaigns leading to a vast number of data-rich images. Currently, to analyze data of this scale, available open-source software (*i.e.*, CellProfiler) requires computational resources that are not available to most laboratories worldwide. In addition, the image-embedded cell-to-cell variation of responses within a population, while collected and analyzed, is usually averaged and unused. Here we introduce SPACe (<u>S</u>wift <u>P</u>henotypic <u>A</u>nalysis of <u>Ce</u>lls), an open source, Python-based platform for the analysis of single cell image-based morphological profiles produced by CP experiments. SPACe can process a typical dataset approximately ten times faster than CellProfiler on common desktop computers without loss in mechanism of action (MOA) recognition accuracy. It also computes directional distribution-based distances (Earth Mover’s Distance – EMD) of morphological features for quality control and hit calling. We highlight several advantages of SPACe analysis on CP assays, including reproducibility across multiple biological replicates, easy applicability to multiple (∼20) cell lines, sensitivity to variable cell-to-cell responses, and biological interpretability to explain image-based features. We ultimately illustrate the advantages of SPACe in a screening campaign of cell metabolism small molecule inhibitors which we performed in seven cell lines to highlight the importance of testing perturbations across models.

## INTRODUCTION

Measuring biological complexity of physiological and pathological states from single cells to whole organisms is the basis for developing models and analytical methods that result in new knowledge moving towards new interventions. Arguably, one of the most successful attempts in measuring cell states, characterizing biological pathways, and testing thousands of small molecules for drug development has been the application of Cell Painting (CP) or its multiple variants (1–7). CP combines several fluorescent dyes to illuminate cellular structures, allowing for an inexpensive, high content (HC), high throughput (HT) microscopy-based assay that is easily scalable to whole genome knock-out/overexpression and large small molecule inhibitor libraries (8–12). Information extraction from CP images involves two main steps: firstly, identifying regions of interest (e.g., cellular substructures) and extracting relevant features, typically done using open-source (*i.e.,* CellProfiler, (13,14)) or commercial software; secondly, performing feature reduction and representation (1,15–17) to facilitate further downstream analyses such as clustering and classification. Due to its biological and analytical relevance, single cell data in HT imaging-based campaigns is now widely used both for data quality control and for hit identification associated with various treatments based upon clear phenotypic differences (6,18–23). Nonetheless, while CP approaches have entered the mainstream for phenotypic screening, there is still a very active research effort to enhance robustness, processing speed, and sensitivity to best capture cell population heterogeneity. As AI-driven strategies have become more prevalent for high-dimensional and large-scale data analysis, integration of such strategies into CP and its variants has stimulated a major interest due to the promise of higher accuracy, faster processing times and the potential for fusing multimodal data (18,19). However, with the continual advancement of HT microscopes and laboratory automation, and resultant large datasets, a key roadblock appears to be how to analyze data efficiently, in a timely manner, and the availability of massive computational resources needed to carry out such analysis.

To address these outstanding challenges, we developed an open-source, Python-based, easy-to-deploy, single-cell image analysis platform named SPACe (<u>S</u>wift <u>P</u>henotypic <u>A</u>nalysis of <u>Ce</u>lls), that tackles object segmentation, quality control, feature extraction and analysis of large image datasets collected from HC/HT imaging campaigns while requiring significantly less computational resources with respect to existing methods. In fact, SPACe can process large datasets commonly used in HT imaging-based campaigns approximately ten times faster than CellProfiler using a standard personal computer, without performance loss in downstream analysis (*e.g.,* MOA recognition accuracy).

The novel SPACe platform includes additional properties that were designed to ensure reproducibility and the ability to improve interpretability of downstream analysis of extracted features. Specifically, SPACe includes a state-of-the-art approach for cell segmentation, the ability to segment multiple subcellular compartments and the application of a directional Earth Mover’s Distance (EMD) variation to quantify differences in single cell feature distributions. We recall that most of existing cell screening analytical platforms, while extracting information from individual cells, ultimately utilize only per well or per treatment central tendency values like mean or median for downstream analysis. A notable exception is the work from Pearson et al., (18) that demonstrated the potential advantage of interrogating single cell information by analyzing the statistical distribution of data within cellular populations. While the traditional per-well average approach has proven to be successful and sufficient for hit calling from single end point assays in large scale screening campaigns, it ignores the inherent phenotypic heterogeneity in a cell population and the fact that many biological responses do not follow a normal distribution (6,19,23).

While the SPACe platform is applicable to virtually any screening campaign using various instruments, we focus here on analyzing images captured with the JUMP Consortium data acquired with vetted high throughput widefield and spinning disk confocal microscopes.

## RESULTS

### Development of a single cell-based image analysis pipeline – SPACe

The traditional way of analyzing CP images relies on either the open-source CellProfiler (1,13) or available commercial software (*i.e.,* Columbus and others). For the larger datasets typically associated with screening, it is recommended to run these solutions on distributed computing resources (*i.e.*, CPU clusters), Cloud computing, or powerful workstations. However, many labs around the world considering using CP may not have access to such resources. To overcome this limitation, we developed SPACe (**Figure 1A-B**), an open-source, user-friendly Python-based analysis pipeline that can efficiently analyze CP images using a standard desktop PC equipped with a consumer grade graphics processing unit (GPU). SPACe was measured to be approximately 10x faster than the open source CellProfiler pipeline (**Figure 1C**), and is provided to the community on Github (https://github.com/dlabate/SPACe) as a downloadable version for local installation, customization, and possible linkage to additional user-defined modules. We also include a Google Colab version of the code (https://github.com/dlabate/SPACe/blob/main/SPACe_colab.ipynb), designed for testing the software under different hyperparameter settings.

**Figure 1.**
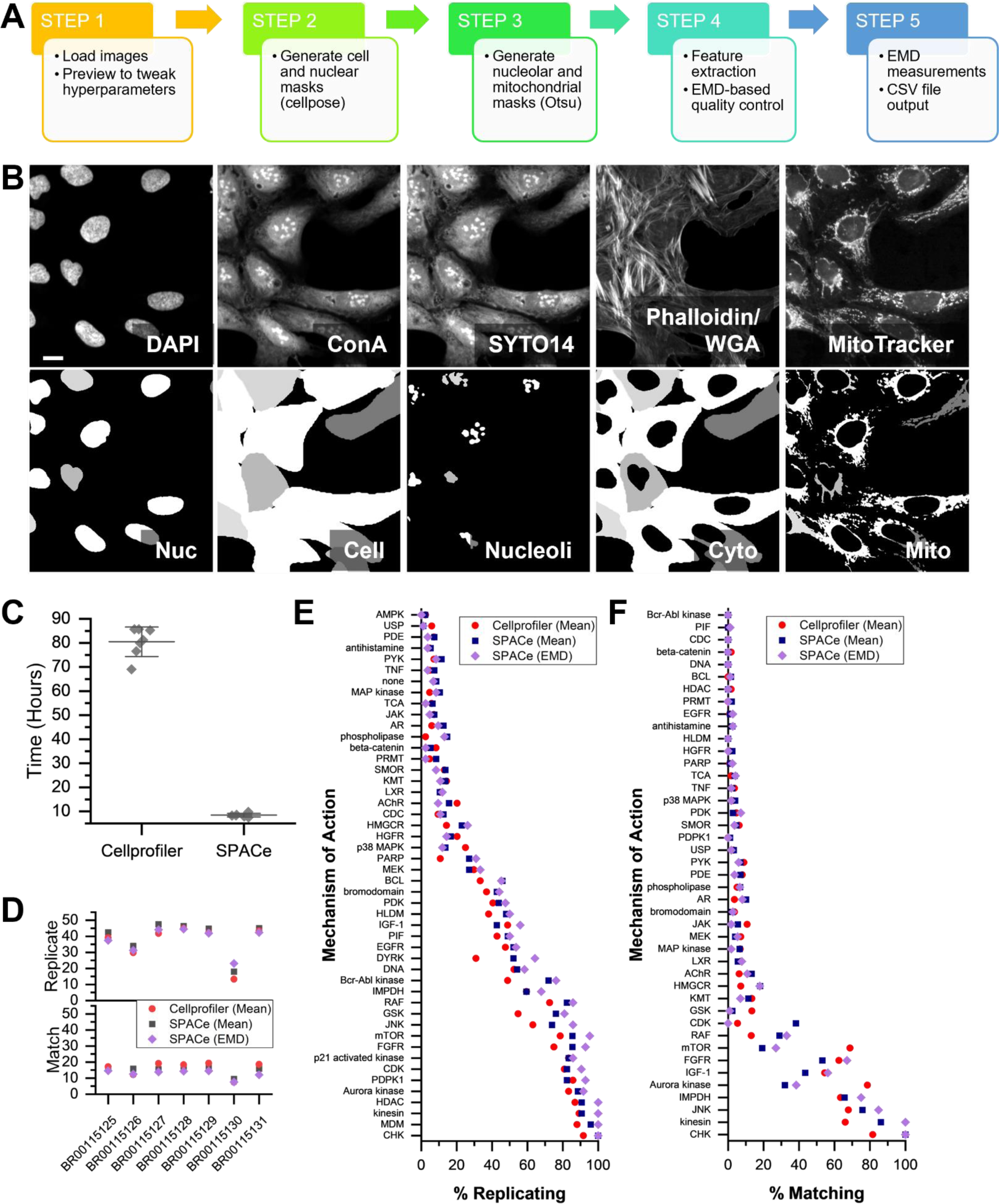
SPACe workflow and performance on JUMP MOA reference datasets. A) A graphical depiction of the steps included in the SPACe pipeline. B) Example of four SPACe segmented objects in CP stained U2OS cells, please note that the fifth compartment (cytoplasm) is obtained by subtraction of the nuclear mask from the cell mask. Scale bar: 5µm. C) Processing time for JUMP MOA reference datasets using either CellProfiler or SPACe. D) Assessment of percent replicating (top) and percent replicating (bottom) based on population mean (CellProfiler, SPACe) or EMD (SPACe only). Assessment of analysis feature percent replicating (E) or percent matching (F) by the mechanisms of actions present in the JUMP reference datasets.

As part of streamlining the CP approach, we decided to not include a module for illumination correction due to the fact that this can be performed as a preprocessing step following established methods (2,24,25). We have often found that instrument-associated image correction is sufficient to remove illumination errors. In the case of experiments performed in this study, we relied upon the Yokogawa CV8000 software to correct non-uniform illumination, pixel misalignment between cameras, and fluorescence channel crosstalk.

After loading images (Step 1), the SPACe pipeline automatically defines nuclear (“Nucleus”) and cell (“Cell”) segmentation boundaries using the popular AI-based Cellpose package and its pretrained generalist model(s) (26,27) (Step 2). Although Cellpose can automatically determine segmentation hyperparameters on a per image basis, for more consistent and faster performance, users should indicate the expected nuclear and cell diameters in terms of number of pixels, which is often cell model specific. To facilitate this task in SPACe, we implemented a preview function that allows the user to test selected fields of view (FOVs) making sure that the segmentation is accurate before analyzing the entire dataset. Following nuclear and cell regions identification, an adaptive Otsu & MaxEntropy thresholding routine is applied to identify nucleoli (“Nucleoli”) and mitochondria (“Mito”, Step 3). A separate cytoplasmic region (“Cyto”) is defined by subtracting each nuclear region from each cell region. An example of segmented objects is shown in **Figure 1B**.

SPACe extracts more than 400 curated features from each object mask including intensity, morphology, and textural measurements (Step 4) that are described in **Table S1**. Again, experienced users can alter/add/subtract the set of extracted features as needed. As we describe in detail below (**Figures S2-S3**), we demonstrate that the number of features extracted by SPACe is sufficient to capture phenotypic changes in Mechanism of Action (MOA) plates produced by the JUMP consortium.

A version of our previously published (19) quality control (QC) pipeline was added in SPACe (Step 4); this step leverages the analysis of the empirical probability distribution of single cell features in control samples (*i.e.,* DMSO treated) to identify and discard outlier wells and to generate a reference distribution for each feature, defined as the median empirical distribution of all remaining DMSO wells. This reference distribution is then used to quantify the effect of perturbations based on the Earth Mover’s Distance (EMD), a metric that quantifies the dissimilarity between probability distributions and has been shown to work well for phenotypic screens (18). Additional details, including data normalization, can be found in the Methods section. Here we adopt a directional variant of the EMD, where we assign a positive sign to the EMD if the median has increased with respect to DMSO, and a negative sign if the median has decreased. The SPACe pipeline ultimately provides users with the saved object masks, single cell raw data, and a CSV file containing the EMD values plus the well-averaged mean and median values for each feature (“distance maps” – Step 5); all of which can be further analyzed using preferred software packages as suggested in several publications (1,15–17). Filtering wells with a low number of detected objects can also be used as an additional QC metric to avoid analyzing the distribution of single cell data from wells with an insufficient number of data points. In our previous study (19), albeit not using CP, we estimated that a minimum of ∼1000 cells are needed to properly reconstruct a faithful distribution from single cell data.

### Comparing SPACe with CellProfiler using JUMP Consortium reference datasets

As one of the goals of SPACe is to be useful to a large community of researchers with potentially limited computational resources availability, we compared its performance to CellProfiler using a highly diverse CP experiment by downloading seven reference datasets provided by the JUMP consortium. These JUMP-MOA datasets (BR00115125-31) contain 90 unique treatments with 47 annotated mechanisms of action (MOA) along with negative control DMSO wells (1). Using a standard PC, of the type available to most research labs (Intel i7 13700 CPU, NVIDIA 3070 GPU, 32GB memory), the average processing time per plate was almost 10x lower using SPACE (8.5 ± 0.5 h) compared to CellProfiler running the pipeline recommended by the JUMP consortium (80.2 ± 5.3 h) (**Figure 1C**). When using the percent replicating and percent matching (1) calculations that define the correlation of extracted features between replicate wells and those with different treatments, but matching annotated MOAs, there was no significant difference between SPACe and CellProfiler generated outputs (**Figure 1D**), with the largest determinant in performance being the source dataset. However, when percent replicating was ranked for each dataset, SPACe mean well values ranked significantly better (**Figure S1A-B**). In percent matching, both SPACe and CellProfiler mean well values ranked significantly better than SPACe EMD values (**Figure S1C-D**). When examined by individual MOAs, the trend in percent replicating and percent matching were similar between analysis methods (**Figure 1E-F**). Although no difference was statistically significant, there were examples (*e.g.,* GSK, Bcr-Abl kinase, Aurora kinase, mTOR) where one method appeared to outperform the other, likely reflecting differences in the underlying segmentation approaches used.

One of the key differences between CellProfiler and SPACe is the reduced number of features extracted (∼4,000 versus ∼400). To understand the uniqueness of the features collected by each method, Spearman correlation between features across all samples in the JUMP reference datasets were examined (**Figure S2A-C**). A large fraction of features is highly correlated (>0.8), regardless of the analysis method being used. The distribution of correlation values showed an enrichment of CellProfiler features below 0.2 and SPACe, for either well mean or EMD values, above 0.8 (**Figure S2E**). When unique features, defined as those with no correlation with another feature above 0.95, were counted, SPACe produced 101.0 ± 6.0 well mean and 103.6 ± 9.5 EMD features that were considered unique (**Figure S2D**). Although CellProfiler produced a larger number of unique features per analysis (563.0 ± 9.0), this represented a smaller percentage (16%) of the total features set than with either SPACe well mean (24%) or SPACe EMD (32%) analysis. This suggests that although SPACe collects a smaller feature set, the feature set contains both redundancy and uniqueness sufficient to recapitulate the CellProfiler generated results from the reference dataset, similar to other published work that reduced the CellProfiler feature set to a little over 600 (28).

A concern with collecting a smaller number of features is the potential inability to accurately capture phenotypes associated with various MOAs. To address this concern, we generated five random forest (RF) models for each JUMP reference dataset for each of the CellProfiler well mean, SPACe well mean, and SPACe EMD feature sets. Each RF model was trained with half of the treatment replicates, randomly selected for each model replicate. Model performance was assessed by the ability to correctly predict the MOA of treatment replicates not selected for training. Across all samples, the feature set used to train the model did not make a significant difference in the accuracy of MOA prediction (**Figure S3A**). When examined by individual MOA, RF models trained with the different feature sets showed similar trend in prediction accuracy (**Figure S3B**). RF models trained with SPACe (either well mean or EMD) features were noted to better predict several MOAs, particularly the DYRK (significant, p < 0.05), MEK, LXR, PRMT, and beta-catenin signaling pathways. When examined by prediction accuracy rank for each MOA, RF models trained with SPACe EMD values ranked significantly better (**Figure S3C**). When a confusion matrix was examined for each set of RF models (**Figure S3D-F**), all incorrect MOA predictions shared a similar pattern, which was to predict a sample as inactive vs. active (*i.e.* “none” category in the graphs). Taken together, these results indicate that the reduced feature set collected by SPACe, especially the EMD-based feature set, is able to capture induced phenotypes as well as, if not slightly better, than the feature sets generated using the JUMP consortium CellProfiler pipeline.

### Testing selected reference chemicals with the pipeline and their reproducibility

We used the standard JUMP consortium U2OS osteosarcoma cell line (1–3) to investigate reproducibility and interpretability of the results obtained with SPACe. We chose a selection of potential reference chemicals (**Table 1**) that have been previously shown in publications or by the JUMP consortium to alter the phenotype of U2OS cells (3,29,30), including saccharin and sorbitol as negative controls. Cells were treated for 24 hours ahead of the CP protocol, which was performed either manually or robotically following published reports (1–3) and imaged using a Yokogawa CV8000 high throughput spinning disk confocal using the established conditions (31). **Figure 2A** shows representative color images of the indicated compounds. In **Figure 2B** the EMD from the median DMSO control (which has distance=0) for each of the >400 features, from three independent biological replicates (labeled 1,2,3 in parentheses below the treatment names), is shown as a heatmap for the “reference compound set” with each of the fluorescence channels separately highlighted. The quantification of all the data in **Figure 2B** is shown in **Figure 2C**, where we used Euclidean distance between the treatment fingerprint (*i.e.,* represented by changes in all >400 features) for each well (represented as a circle in the graph), from the median of DMSO wells of each biological replicate. Overall, in terms of Euclidean distance, we observed good concordance between the three independent biological experiments, and a compound with Euclidean Distance >2 was considered active as it was fully separated from the DMSO control wells and was significant by non-parametric ANOVA, a parameter that we used for the rest of the study. As expected, the negative controls (saccharin and sorbitol) had no effect across the feature space. Of the tested reference compounds, we also failed to detect significant and/or reproducible effects of tetrandrine, dexamethasone and NVS-PAK1-1. Berberine chloride caused a redistribution of well-resolved mitochondria, which was evident from the images (**Figure 2A**) and was readily measured by changes exclusively in the MitoTracker channel (**Figure 2B** and **2D**). To aid with interpretability of the results (**Figure 2D**), we subdivided the features in categories (based on cell compartment and fluorescence channel) and then represented changed features (defined as EMD >0.15 or <0.15 in at least two out of three biological replicates) as a stacked bar graph (blue features are reduced and red features are increased as compared to DMSO). AMG-900 and Etoposide showed changes in the nuclear and nucleolar compartments, which are compatible with their known mechanism of action (aurora kinase and topoisomerase inhibitors, respectively). Rotenone affected multiple compartments, including mitochondria. Similarly, fenbendazole and oxibendazole, two anti-parasitic drugs, showed changes across all compartments, including visual evidence of polynucleation, as previously described (3). In the case of Ca-074-Me, we did not observe the increase in ConA staining intensity that was previously reported (3), but the most reproducible changes were measured at the level of nuclear and cellular size and shape. TC-S-7004, a DYRK1A/1B inhibitor (32), also had a fingerprint that was significantly different from DMSO, with novel changes in the SYTO14 and MitoTracker channels.

**Figure 2.**
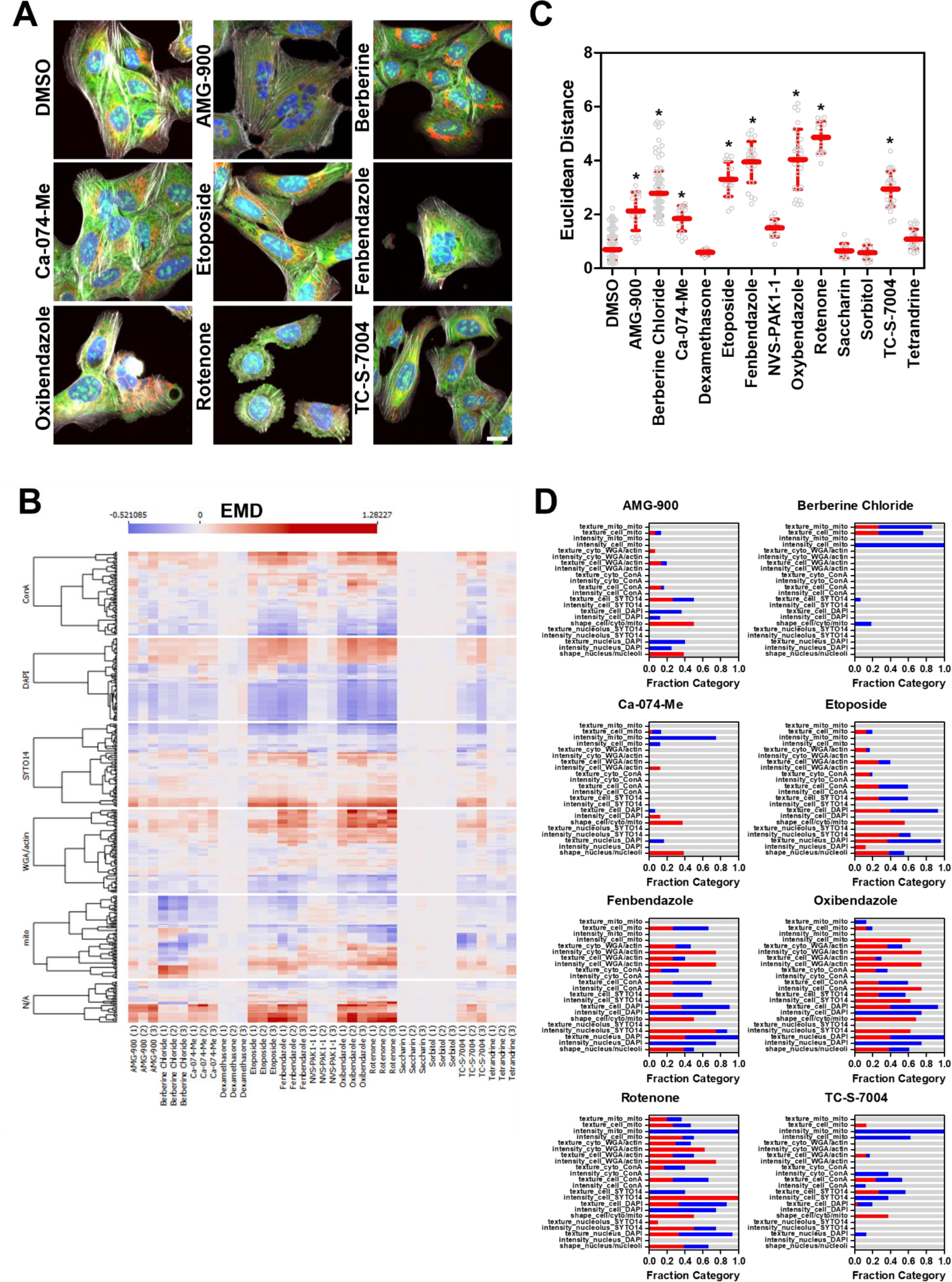
Testing a set of potential reference compounds with SPACe. U2OS were treated for 24 hours with the indicated compounds at 10µM ahead of the CP protocol. Representative 20x color images are shown in panel A. Scale bar is 10µm. B) Heatmap showing the EMD from DMSO controls of the indicated compounds from three independent biological replicates (labeled in parentheses below the name of the compounds as 1,2,3). Each channel is separated to show the changes by fluorescent dye. C) Euclidean distance is used as a measure to compare the features fingerprint of each compound from the median DMSO control. This is measured for each well (represented as a hollow circle in the graph) treated with the indicated compound in three independent biological replicates. * is p<0.01 using non-parametric ANOVA compared to DMSO group. D) stacked-bar graphs representing the fraction of changed features (in blue if they are reduced or red if they are increased, grey means no change) in the defined groups for each treatment. The chosen threshold for significance was EMD distance <-0.15 or >0.15 (see Methods for description).

**Table 1.**

| Chemical Name | CAS # | Literature Reported Phenotype | Notes |
| --- | --- | --- | --- |
| AMG-900 | 945595-80-2 | JUMP positive control | Aurora kinase inhibitor |
| Berberine Chloride | 633-65-8 | Redistribution of mitochondria | Target unclear |
| Ca-074-Me | 147859-80-1 | Bright ConA staining | Cathepsin B inhibitor |
| Dexamethasone | 50-02-2 | JUMP positive control | GR agonist |
| Etoposide | 33419-42-0 | Large nucleoli | Topoisomerase II inhibitor |
| Fenbendazole | 43210-67-9 | Multi-nucleated cells | Anti-helminthic |
| NVS-PAK1-1 | 1783816-74-9 | JUMP positive control | PAK1 inhibitor |
| Oxibendazole | 20559-55-1 | Multi-nucleated cells | Anti-helminthic |
| Rotenone | 83-79-4 | Mitochondrial fission | Complex I inhibitor |
| Saccharin | 81-07-2 | Negative control | Artificial sweetener |
| Sorbitol | 50-70-4 | Negative control | Sugar alcohol |
| TC-S-7004 | 1386979-55-0 | JUMP positive control | DYRK1A/1B inhibitor |
| Tetrandrine | 518-34-3 | Abundant WGA | Calcium channel blocker |

### Reproducibility and interpretability: Berberine chloride as a case study

To further analyze the quality of SPACe pipeline outputs, we first focused on berberine chloride as it showed a unique phenotype that is immediately evident from visual inspection of the images (*e.g.,* redistribution and size reduction of mitochondria). In **Figure 3A**, the analysis of thirteen independent biological replicate experiments confirmed high reproducibility of the berberine-induced, visible phenotype in terms of the mitochondria features changing from the DMSO control. Only one experiment was somewhat an outlier with additional cellular compartments being more strongly affected; however, the top mitochondrial features remained similarly altered. We performed berberine chloride dose-response experiments (four biological replicates) to verify if the top changing mitochondrial features were indeed responding in a concentration-dependent manner. **Figure 3B** shows the berberine chloride dose-response experiments in a heatmap format, confirming that most responding features indeed have a dose dependency, an indication of their biological specificity likely linked to the compound’s mechanism of action. We next selected eight features (intensity, texture, plus the ratio between mitochondrial mask and cell mask areas, **Figure 3C)**, that exhibit clear dose dependency, with very similar EC_50_s, indicating they are likely related to the berberine chloride mechanism of action, which remains ill-defined (33,34).

**Figure 3.**
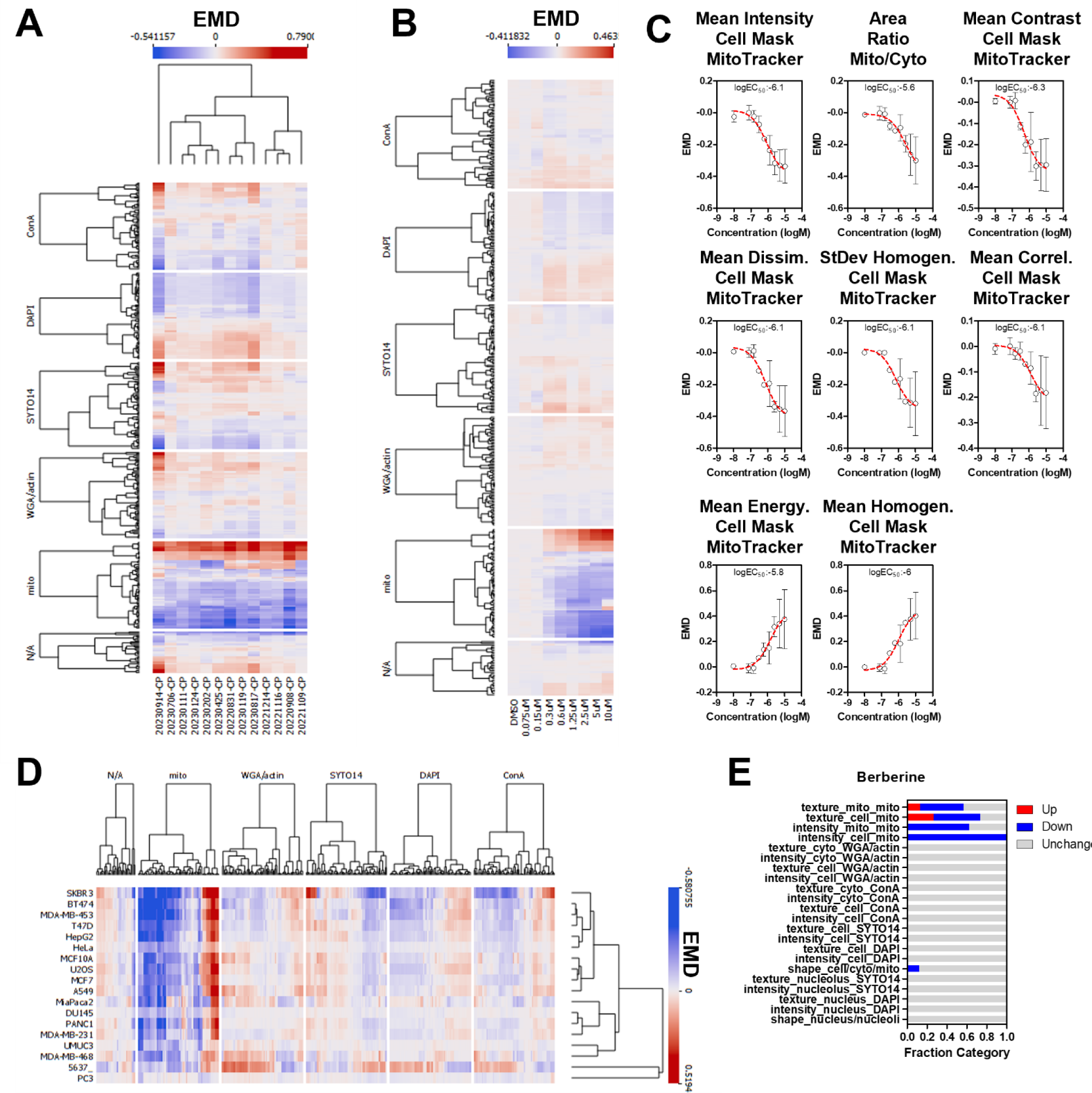
Berberine chloride as an example of a reference compound with a clear, interpretable phenotype. A) Feature response reproducibility. Thirteen independent experiments were conducted with U2OS cells treated with berberine chloride (10µM) for 24h, features were extracted with SPACe and EMD is represented as a heatmap. B) EMD heatmap showing all the extracted features after a berberine chloride dose-response (75nM to 10µM) at the 24h time point. C) average +/- standard deviation of the top, non-redundant eight features changing after berberine chloride treatment, with EC_50_ indicated. D) eighteen cell lines were treated with berberine chloride 10µM for 24 hours and the EMD of all features is represented as a heatmap. E) stacked bar graph showing a berberine chloride “consensus fingerprint” of the changing features across cell lines.

To determine if the response to berberine chloride was universal so that it could be employed as a true reference compound for all CP experiments, we treated 18 human cell lines for 24 hours (**Figure 3D**). 14 out of 18 cell lines showed a strong mitochondrial phenotype, while three more had a partial responsiveness to berberine chloride, reinforcing the assumption that the mechanism of action of this chemical is largely universal and visually affects primarily mitochondria. Moreover, it is interesting to point out that berberine was non-toxic in all the cell lines tested. PC3, a prostate cancer cell line, was the only clear outlier for unknown reasons, with bladder cell lines UMUC3 and 5637 also having a muted and more diverse fingerprint. The consistent response allowed us to extract a “berberine chloride consensus fingerprint” that is visible as aggregate results in a stacked bar plot subdivided by feature classes and cellular compartments (**Figure 3E)**. A list of the selected features that constitute such fingerprint is available in **Table S2**. Collectively, the fingerprint signature allowed us to add interpretability to this treatment as the selected features can be visually linked to the images. For example, the intensity of the MitoTracker channel in the cell compartment is reduced as is the ratio of mitochondria-to-cell area, while several distinct mitochondrial texture features are changing both within the cell and mitochondria compartments.

### Cell Painting in breast cancer cell lines: analysis of a small panel of chemicals reveals both cell-type specific and broad effects

To further explore the potential of SPACe for analysis of cell lines outside the canonical U2OS and A549 models often used in CP, we tested a set of breast cancer models representing different tumor subtypes (luminal, Her2, and triple negative), by treating them for 24h with a small set of 28 diverse chemicals, including those deemed active in **Figure 2C**. **Figure 4A** shows the Euclidean distance of each compound from DMSO across multiple experiments and cell lines, represented as a heatmap. Overall, 19 out of 28 compounds were found to be active (Euclidean distance >2, 68%) in at least one cell line; however, only 8 out of 28 (29%) were active in at least five cell lines. These included the Akt inhibitor MK-2206, berberine chloride, fenbendazole, oxibendazole, rotenone, TC-S-7004, actinomycin D, and latrunculin B; all of which have been used as references before or have established strong responses and mechanism of action. Interestingly, we had no compound that showed specificity for U2OS, however, a few were only active in one breast cancer cell line, including BYL719 (PI3K inhibitor, BT474), DCA – deoxycholic acid (bile acid, SKBR3), ETP45658 (PI3K inhibitor, BT474), FR180294 (ERK1/2 inhibitor, MDA-MB-231), and metoclopramide (dopamine receptor antagonist, MDA-MB-231). A more detailed inspection of the phenotypic changes revealed that the two PI3K inhibitors had a very similar overall profile in BT474, indicating a likely class (and perhaps cell line) specific phenotypic readout (**Figure 4B**). In contrast, the two MDA-MB-231 specific compounds modulated multiple compartments and channels, being nucleolus and actin for metoclopramide, mitochondria and concanavalin A/SYTO14 for FR180294.

**Figure 4.**
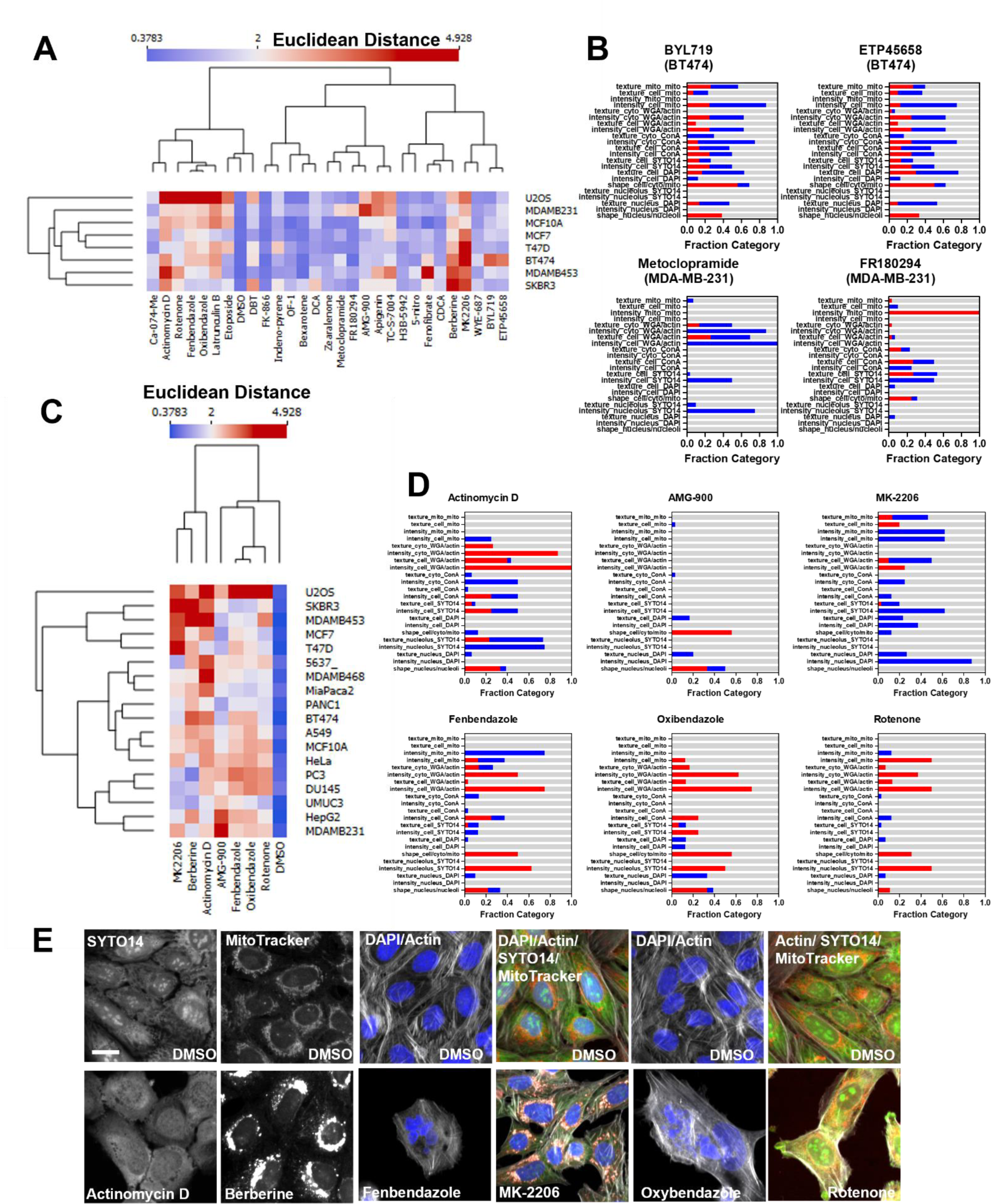
Testing SPACe on a set of breast cancer models. A) Cell lines were treated with 28 chemicals for 24h and then analyzed with SPACe. The heatmap shows the Euclidean distance of each compound from the DMSO wells of each cell line. B) stacked bar graphs showing the interpretability profile of four cell line specific hit chemicals. C) Eighteen cell lines were treated with the indicated compounds for 24 h and then Euclidean distance was calculated and represented as a heatmap. D) stacked bar graphs showing tentative “consensus fingerprints” of the changing features across a minimum of five cell lines. E) selected images (zoom in from 20x/1.0 images) to showcase specific phenotypic changes caused by the indicated compounds in highlighted channels/compartments. Scale bar: 10µm.

We then selected seven of the best responders and tested them across all 18 cell lines (**Figure 4C**) to determine their universality and attempt to identify their response fingerprint, akin to what we showed for berberine chloride in **Figure 3E**. Overall, all compounds elicited a response in at least 10 cell lines, but none in all 18, indicating that it is unlikely to identify compounds that would act in a universal manner and can be used as controls across all experimental models. This complicates the analysis, prediction, and interpretation of the MOA for compounds when based uniquely on phenotypic screening in a single cell model. We attempted to add interpretability by selecting only features that were changing in at least 5 cell lines, as very few to none significantly changed across all models. **Figure 4D** shows the interpretation using stacked bar graphs for all seven compounds, except for berberine chloride that was already presented in **Figure 3E**. Actinomycin D showed major changes in the nucleolus (**Figure 4E**), compatible with its activity as a general inhibitor of RNA polymerases and gene transcription, and actin cytoskeleton, which can represent signs of toxicity, even though we were still able to collect information from more than 1000 cells/experiment. For AMG-900, only a few features were consistent across cell lines, and these revolved around cell and nuclear size/shape and DAPI texture features, which is compatible with its known mechanism of action in mitosis as an Aurora kinase inhibitor. MK-2206 is a specific Akt inhibitor that can be an autophagy activator (35) and the main features that appear to be linked to this phenotype include reduction in nuclear DAPI signal and changes in texture features for SYTO14, MitoTracker, and WGA/phalloidin that can be visually interpreted as the formation of autophagolysosomes in the cytoplasm (**Figure 4E**).

Fenbendazole and oxibendazole are anti-parasitic drugs that have been shown to act through multiple mechanisms including microtubule destabilization, G2/M arrest, and apoptosis (36,37). Both drugs produced complex CP profiles modifying all the compartments in various ways, most notably higher SYTO14 intensity in the nucleolus and reduced MitoTracker signal in mitochondria. These observations highlight the utility of including these subcellular compartment masks in SPACe. Inspecting the images clearly shows that both compounds cause multinucleated cells and dying cells, confirming their likely mechanism of action (*e.g.*, cell division and cell death, **Figure 4E**). Finally, rotenone, a natural isoflavone, is a strong inhibitor of mitochondrial complex I, which is reflected by the alterations observed in selected MitoTracker features. Interestingly, rotenone appears to cause broader changes also affecting the nucleolus and actin cytoskeleton.

### Cellular metabolism screening library

To expand our understanding of the universe of phenotypic changes across models, we treated seven cancer cell lines with different origins (U2OS - bone, HepG2 - liver, 5637 - bladder, PANC1 - pancreas, PC3 - prostate, MDAMB231 - breast, and A549 - lung) in duplicate plates with the Cayman Chemical Cellular Metabolism Screening Library containing 160 small molecule modulators of diverse targets and metabolic pathways. First, we analyzed the responses to the library by identifying toxic compounds. In **Figure 5A**, treatments that caused a reduction in cell number by >50% in at least one cell line, as compared to DMSO control, are shown in a heatmap format. Overall, about a third of the library showed some toxicity in at least one cell line. Ten compounds were the most toxic across all models (auranofin, SF1670, plumbagin, PR-619, CB-5083, PFK158, eeyarestatin 1, digitoxin, paclitaxel, and TG101348) and should be tested at lower concentrations to measure changes at non-toxic levels, as some of them show potentially very interesting phenotypes in the surviving cells. For example, PR-619 and eeyarestatin-1, in U2OS and across cell lines, respectively, show cytoplasmic vacuolization and redistribution of mitochondria; plumbagin, in PANC1 cells, also affects mitochondria plus on cell shape/size; while TG101348, in PANC1 and HepG2, causes large changes in the WGA/actin, MitoTracker and some morphological features (**Figure S5**). Examples of compounds that showed some cell type selective toxicity were gemcitabine (PC3, nucleoside analog), copanlisib (5637, PI3K), mycophenolic acid (5637, inosine-5’-monophosphate dehydrogenase), and NCT-503 (HepG2, PHGDH).

**Figure 5.**
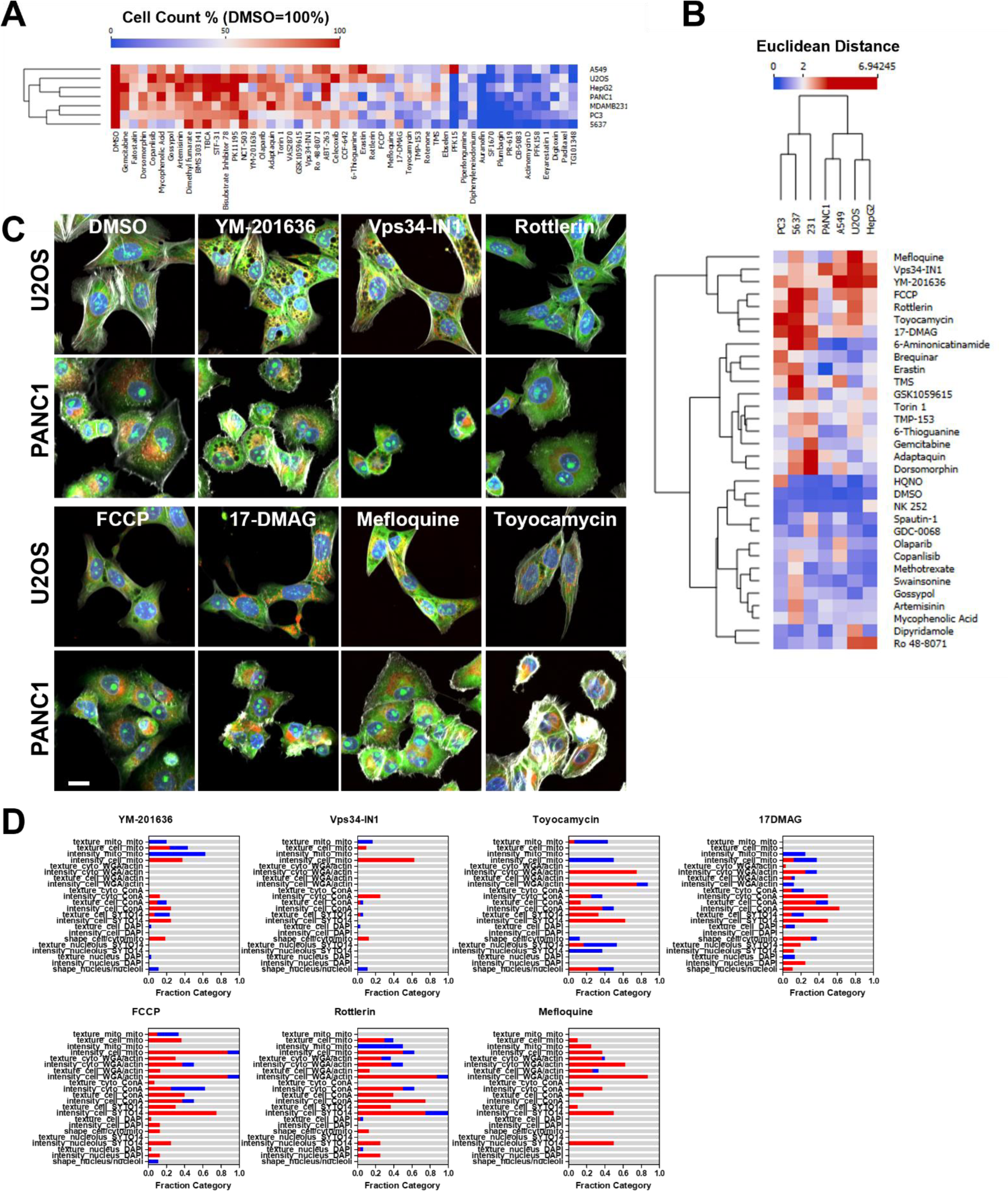
Screening a cellular metabolism modulators library with SPACe analysis. A) analysis of the library compounds induced toxicity represented as heatmap where cell count was normalized to DMSO, set as 1; the compounds represented caused a cell loss of>50% in at least one cell line. B) heatmap and hierarchical clustering of hits (Euclidean distance >2 in at least one cell line) from the screen after SPACe analysis. C) 20x zoomed in images of the indicated hits shown in U2OS and PANC1 cells. Scale bar: 10µm D) Interpretability stacked bar graphs for the compounds shown in panel C.

In **Figure 5B** we present the 31 hit compounds from the library screen in a heatmap, defined as treatments with Euclidean distance>2 cutoff in at least one cell line, and in both replicate plates, after filtering out the abovementioned toxic chemicals. Compounds with discordant replicates or with obvious imaging artifacts were also excluded after manual inspection. To improve accuracy and interpretability, it is important to note that every multiwell plate was run with several internal controls (DMSO, actinomycin D, MK-2206, and berberine chloride). Additionally, actinomycin D and rotenone were present as components of the library itself and served as additional quality control treatments as active compounds, and as such were excluded from **Figure 5B**.

Perhaps interestingly, only seven compounds in the screed showed effects in a single cell line: HQNO (PC3), mycophenolic acid (5637), olaparib (A549), GDC-0068 (MDA-MB-231), dipyridamole (U2OS), spautin-1 (MDA-MB-231), and NK 252 (HepG2); further validating the importance of screening across a wide range of cellular models. Seven compounds showed various phenotypes across four or more cell lines, despite having a somewhat reduced cell number in specific models (top cluster in **Figure 5B**). These were the PIKfyve inhibitor YM-201636, the VPS34 inhibitor Vps34-IN1, the mitochondrial uncouplers FCCP and rottlerin, the IRE1 inhibitor toyocamycin, the Hsp90 inhibitor 17-DMAG, and the antimalarial mefloquine. Visual representations of the phenotypes induced by these compounds are shown for U2OS and PANC1 cells in **Figure 5C**, while their interpretability plots are shown in **Figure 5D**. In the case of all these compounds, it was much harder to identify a common fingerprint between cell lines, which was especially true for Vps34-IN1, where only 25 features were common between three out of seven cell lines. However, in individual cell lines, visually, YM-201636 and Vps34-IN1 responses looked reasonably similar (U2OS cells are shown in **Figure 5C**), and indeed we found a set of features that matched these two treatments. For example, increased intensity of the concanavalin A and MitoTracker in the cell mask, reduction in nuclear perimeter and various changes of MitoTracker textural features. Also, the two mitochondrial uncouplers, rottlerin and FCCP, had several matching features in the mitochondrial compartment (*e.g.,* increase intensity and mitochondria/cell area ratio).

## DISCUSSION

Over the last few years, the leading approaches to HT genetic and chemical screens have shifted from classical single-endpoint cell free assays to unbiased multi-endpoint assays based on imaging phenotypic profiling of intact cells. The success of the latter approach stemmed from the development of CP protocols that allow for economical and fast characterization of a cell state by illuminating specific cellular components. This strategy, coupled with automated image analysis and machine learning, has been proven to be very effective in measuring phenotypic changes upon a large variety of perturbations, including small molecules, knock-downs and over-expression.

A major limitation to a wider adoption and deployment of CP-like phenotypic screening is the significant computational resources required to run current image analysis solutions such as CellProfiler and other commercial applications. To address this challenge, here we introduce SPACe, an open-source, lightweight, Python-based CP analysis workflow that differs from most current analysis tools in several important ways. First, SPACe integrates the use of Cellpose pre-trained AI-based nuclear and cell segmentation models to achieve high accuracy object segmentation accelerated by widely available GPU-based processing; moreover, we segment two additional cell compartments, *i.e.,* nucleoli and mitochondria, to provide more specific biological information and improve interpretability of downstream analysis. Second, SPACe collects a carefully selected set of ∼400 image-based features as compared to the ∼4,000 features collected by CellProfiler, reducing redundancy and data management loads; as we have shown, this implementation choice does not reduce in any way the downstream analysis performance while it makes data exploration more efficient. Third, SPACe includes the calculation of directional EMD for feature analysis, as EMD values have been shown to be superior (18) in capturing heterogeneity of responses in the cell population and can be used for both quality control and hit calling (19,21–23), while providing the canonical per well statistics of other platforms.

Due to these design properties, the SPACe pipeline is able to analyze the large imaging data generated by typical CP phenotypic screens very efficiently using the computational resources of a standard PC while maintaining sufficient morphological sensitivity and specificity to train predictive machine learning models for treatment targeted MOAs that perform as well, or better, than predictive models derived from much larger feature sets. This confirms the competitivity of our feature selection process given that MOA prediction is known to be a very challenging task (15,38,39).

The ability of SPACe to efficiently capture single-cell morphological features also explains the potential of this pipeline for providing biological interpretation of image-based fingerprints. In this study, we applied SPACe to define phenotypic fingerprints of common reference compounds, small targeted chemical sets, and a larger chemical library targeting cellular metabolism in U2OS cells and then expanding up to 18 different cell models. We demonstrated that very often, potent well-defined reference compounds do elicit a phenotypic response across most (but never all) cell lines. However, we found that changes in the underlying features are rarely the same, making it very challenging to generate “universal fingerprints” that could be used to establish universal reference compounds. In larger screens, most chemicals elicit cell-type and feature specific effects. This is not unexpected due to each cell model representing a unique set of activated cellular signaling pathways upon which the chemical perturbation may alter. This is one reason why it is difficult to train a single prediction model to identify and predict MOA-specific phenotypes that are accurate across multiple cell models. However, exceptions exist, the most prominent being berberine chloride which affects only the mitochondrial compartment across almost all models we tested. This finding allowed us to extensively test and conclude that berberine chloride-induced phenotypic changes and the SPACe-extracted features are indeed highly reproducible across multiple biological replicates performed months apart, and across multiple cell models. Further work is needed to determine if this phenomenon is specific to berberine chloride and the mitochondrial compartment, or if other compartment-specific perturbations could be identified.

In conclusion, SPACe offers an open source, user-friendly and highly efficient platform for the analysis of single-cells HT and HC phenotypic screenings. Due to its lightweight implementation, we expect that this computational software will be particularly beneficial to the large community of researchers who are interested in exploring CP analysis but do not have access to large computational resources, *e.g.,* multi-core computing clusters, required to run current software solutions (*e.g.*, CellProfiler) to process CP-generated feature data.

## MATERIALS AND METHODS

### Cell culture and treatments

All cell lines were obtained directly from ATCC or from the BCM cell culture core (Department of Molecular and Cellular Biology) and grown in their ATCC suggested media. Cells were routinely checked for mycoplasma contamination by DAPI staining and high magnification imaging. Cells were plated in 384 well optical bottom plates (PerkinElmer cat# 6057302) at a density of 2000-3000 cells/well and allowed to settle at room temperature for 30 minutes prior to being placed in the incubator at 37C/5% CO2. After 24h, without changing media, cells were then treated with the indicated compounds at indicated doses for an additional 24 hours. All chemicals were reconstituted in DMSO at a stock concentration of 20mM, which is 2000x of the highest concentration tested, unless otherwise specified. The basic protocol requires plating cells in 20µl of media, treatments are then added on top in 20µl of media (final dilution of the chemicals is 1000x).

The cellular metabolism screening library (Cayman Chemical, cat #33705, batch #0609421) contains 160 known modulators of metabolic pathways. The library was arrayed in the test plates using a Labcyte Echo 550 acoustic liquid handler at the Texas A&M Institute of Bioscience and Technology, together with DMSO, berberine chloride, actinomycin D, and MK-2206, which were used as reference treatments (negative and positive). A few wells were also included that were not treated to confirm the control DMSO concentration was non-toxic and did not affect extracted features. Every compound in the library was used at 10µM for 24h of treatment before following the Cell Painting protocol described below.

### Cell Painting (CP) processing and imaging

Following the JUMP consortium CP protocol (1), multiwell plates were processed either manually or robotically (Beckman Coulter Biomek i5). In practice, 20µl of MitoTracker (3000x) was added to live cells for 30 minutes and incubated at 37C. 20µl of fixative (16% PFA in PBS – final concentration in the well is 4%) was then added for 15 minutes at room temperature before proceeding with application of the fluorescent dyes. Slight changes were needed for robotic processing, specifically, the treatment media was removed, 20µl of MitoTracker (1000x) was added and, after 30 minutes, 20µl of fixative (8% PFA in PBS, final concentration is 4%) was added.

Plates were imaged inside a three-day window on a Yokogawa CV8000 high throughput spinning disk confocal microscope with sequential imaging of the five CP channels and appropriate laser/emission filter combinations as described in (1,31). Imaging was performed with a 20xW/1.0 objective, and a short z-stack (3x z planes, 1µm apart) was captured to correct for plate unevenness. Images were first processed using Yokogawa software to correct non-uniform illumination, pixel misalignment between cameras and fluorescence channel crosstalk. Max intensity projections were saved as 16bit TIFFS for automated image analysis. A minimum of nine fields of view were collected from each well in an experimental campaign. With these parameters, a single 384 well plate could be imaged in 3 to 5 hours depending on exposure time and number of FOVs with a file size of ∼120Gb. For campaigns with 4 or more plates, imaging was automated using a BioAssemblyBot 400 robot (Advanced Solutions), a fully integrated plate-loading solution synchronized with the Yokogawa CV8000.

### SPACe Image Analysis pipeline details

For downloading and installing SPACe follow this link that contains all the instructions: https://github.com/dlabate/SPACe. Note, for the correct deployment of SPACe, the plate map must be properly formatted (see example provided as **Supplementary Spreadsheet 1** and downloadable at the github link). SPACe was tested on images captured from a Yokogawa CV8000 HT spinning disk confocal, an ImageXpress Micro XLS widefield HT microscope and a Yokogawa CQ1 spinning disk confocal. SPACe includes the following steps (**Figure 1A**):

### Step 1 – load images and select hyperparameters (preview)

After loading the images, the preview function allows users to optimize the hyperparameters linked to identifying the nuclear and cell mask. A Google Colab notebook (https://github.com/dlabate/SPACe/blob/main/SPACe_colab.ipynb) is available for users to preview sample images and set best hyperparameters for segmentation using Cellpose. Hyperparameters include minimum diameter size for nucleus and cytoplasm (default parameters: 100 pixels for both), minimum number of nucleus, cytoplasm, mitochondria, and nucleoli (default parameters: 600, 700, 40, 200, respectively), minimum and maximum of nucleoli size (default parameters 0.005 and 0.3 control lower and upper threshold). Image intensities are rescaled before segmentation to ensure uniform processing across all image channels.

### Step 2-3 – segmentation

This step detects each cell and, within each detected cell, identifies cytoplasm, nucleus, nucleoli, and mitochondria. It also ensures that each cellular subcompartment is assigned to the corresponding cell with the same label. We apply the generalist learning-based segmentation algorithm Cellpose v2.2 (26,27) to the DAPI and ConA channels to generate nucleus, cell, and cytoplasm masks. Cellpose requires a user-defined hyperparameter that estimates expected cell and nuclear diameters in pixels, that should be optimized in Step 1 using the preview function. Following this segmentation step, we apply label matching to ensure that each segmented nucleus and cytoplasm is assigned to the corresponding cell with the same label. This routine corrects potential errors introduced by Cellpose that might mistakenly detect multiple nuclei per cytoplasm, or a cytoplasm without nucleus. Next, we apply another custom-designed segmentation routine based on Otsu and MaxEntropy thresholding to segment nucleoli (using the SYTO14 channel, within the nucleus mask) and mitochondria (using the MitoTracker channel, within the cytoplasmic mask), followed by another label matching step.

### Step 4 - feature extraction and quality control

This step computes single cell features (shape, intensity, and texture) using the five masks described above (cell, cytoplasm, nucleus, nucleolus, and mitochondria) and images from the five acquired channels. We selected ∼400 features that are widely used while paying special attention to features that are biologically interpretable. A complete list is included in **Table S1**.

The QC routine is designed to establish a reliable ground truth for single cell distributions in control samples (*e.g.,* DMSO). The idea stems from our prior publication (19) that demonstrated the value of distribution analysis as a quality control step for high throughput microscopy assays and subsequent single cell analyses. The QC step establishes a reference distribution for the DMSO negative control wells (eliminating outliers because of low object count or aberrant phenotypic profile). The reference distribution is here defined as the median of the DMSO distribution in each experiment. The same QC step can also be applied to each set of replicate treatment wells, if appropriate, to discard outlier wells (*i.e.,* wells with missing/low number of cells, artifacts, no or super response).

### Step 5 – Directional Earth mover’s distance (EMD)

For each well and each feature, this step computes the EMD to the established reference DMSO distribution. Before computing the EMD, features were normalized independently per plate by removing top and bottom 2% and standardized in the interval [-1.1] using the Python command *robustscale* in scikit-learn. A sign (plus or minus) is next assigned to the EMD to indicate the direction of the response as compared to the reference distribution, plus sign indicates that the median value has increased with respect to DMSO and negative sign to indicate that it has decreased.

### SPACe output

The results are available in a set of folders that contain intermediate steps (*i.e.,* masks from each compartment – Step 2 & Step 3 folders), single cell data (Step 4), and distance maps (Step 5). The distance map .csv file contains the EMD values for each analyzed well plus per well mean and median values.

### Analysis of JUMP MOA Reference Datasets

Seven JUMP MOA reference datasets (BR00115125 – 31) were downloaded from the JUMP Cell Painting gallery (https://registry.opendata.aws/cellpainting-gallery/). For analysis time calculations, all datasets were processes using CellProfiler version 4 (**Error! Hyperlink reference not valid.** and SPACe using a desktop PC equipped with a 16-core Intel i7 13700 CPU, NVIDIA 3070 GPU, and 32GB memory. CellProfiler analysis utilized the segmentation and feature extraction pipelines provided by the JUMP consortium (https://github.com/broadinstitute/imaging-platform-pipelines/tree/master/JUMP_production). The illumination correction and quality control pipelines were excluded since equivalent operations are not present in SPACe.

### Calculation of percent replicating and percent matching

Percent replicating and percent matching was evaluated as previously described (1). In practice, single-cell data was aggregated at the per well level by calculating the mean or EMD (SPACe only) value for each feature. All features were then normalized based upon plate Z-score. Using normalized values, we generated a null distribution of 20,000 random non-matching pair-wise correlations (Spearman) for each dataset. Using a threshold based on the 95% percentile of the null distribution, the percentage of replicating or matching well pairs with a correlation above this threshold was determined to calculate the percent replicating and percent matching values for each dataset and/or MOA. All calculations were completed using Biovia Pipeline Pilot (version 18.0) software.

### Generation of Random Forest (RF) MOA prediction models

Data for model generation consisted of mean- or EMD-aggregated and plate Z-score normalized well-values from six JUMP MOA datasets generated using either CellProfiler or SPACe. No data cleaning or handling of missing values was required. For each dataset, replicate wells were evenly and randomly assigned for training or testing purposes. A RF model was initialized with the following hyperparameters: 500 total trees, maximum depth of 50 trees, no minimum samples per leaf, and maximum features per tree set at the square root of total features. The RF model was trained using samples assigned to training. Each decision tree in the forest was constructed using a bootstrapped sample from the training data. At each split in a tree, a random subset of features was considered for splitting. The Gini impurity criterion was used to measure the quality of splits. The trees were grown until they reached maximum depth or when the minimum number of samples per leaf (=1) was met. The trained RF model was used to make MOA predictions of samples assigned to testing. Model performance was evaluated by the percent of correct predictions (accuracy) and a confusion matrix generated. All steps were repeated 5-fold for each dataset and each type of feature data for a total of 90 models. Models were generated using R and its RF library.

### Statistical analysis

All experiments were performed a minimum of three times (biological replicates) with a minimum of 4 wells/treatment (technical replicates), except for the Cell Metabolism screen where only two wells/treatment were used. To compare fingerprints, Euclidean distance was measured between EMDs of all features in the treatment wells and the median DMSO control wells. Euclidean distance is a standard method used to measure the distance between two points in a multi-dimensional space and has been employed from gene expression analysis to image-based morphological profiling (2,5,16,40,41). Groups were compared with non-parametric Kruskal-Wallis test. Heatmaps and clustering were generated using Orange Data Mining v.3.36, graphs were made in GraphPad.

### Interpretation plots

The stacked bar graphs were prepared as follows: features were grouped into classes based on compartment and channel, then, for each treatment, features with EMD>0.15 and EMD<-0.15 were assigned to the class of “changing features” (the threshold was based on the reproducibility analysis of berberine chloride treatment as shown in **Figure 3**). Number of changing features were then transformed into a fraction by dividing over the total number of features (hence, sum=1) and labeled as *up* (EMD>0.15, red in the figures), *down* (EMD<-015, blue in the figures), or *unchanging features* (grey in the figures).

## ACKNOWLEDGMENTS

Software development, experimental approaches and imaging for this project was supported by the Center for Advanced Microscopy and Image Informatics (CAMII, CPRIT RP170719) and the Integrated Microscopy Core at Baylor College of Medicine (funding from NIH (DK56338, CA125123, ES030285, S10OD030414), and CPRIT (RR200043), the Dan L. Duncan Comprehensive Cancer Center, and the John S. Dunn Gulf Coast Consortium for Chemical Genomics.

## SUPPLEMENTARY MATERIALS

**Table S1.**
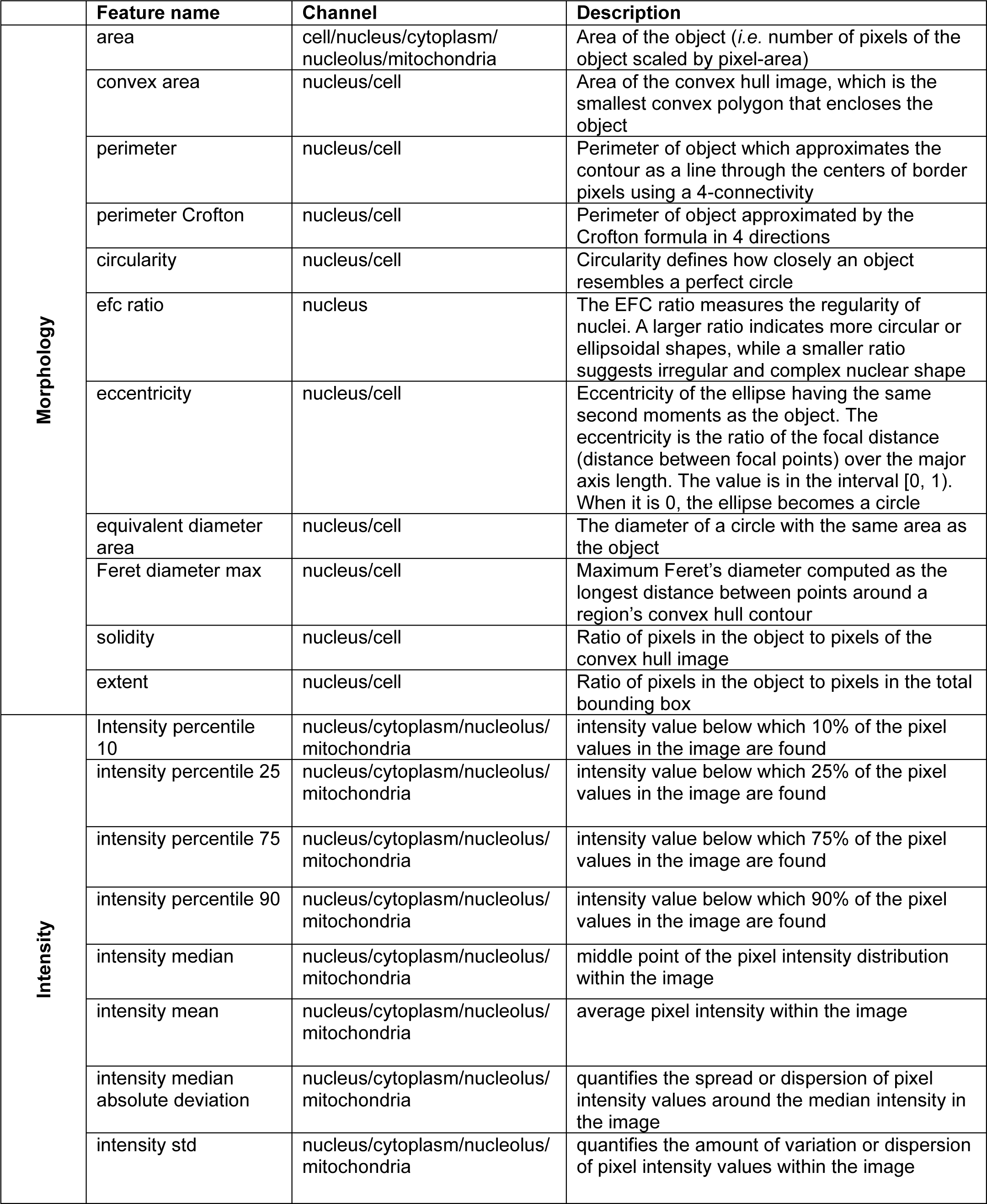
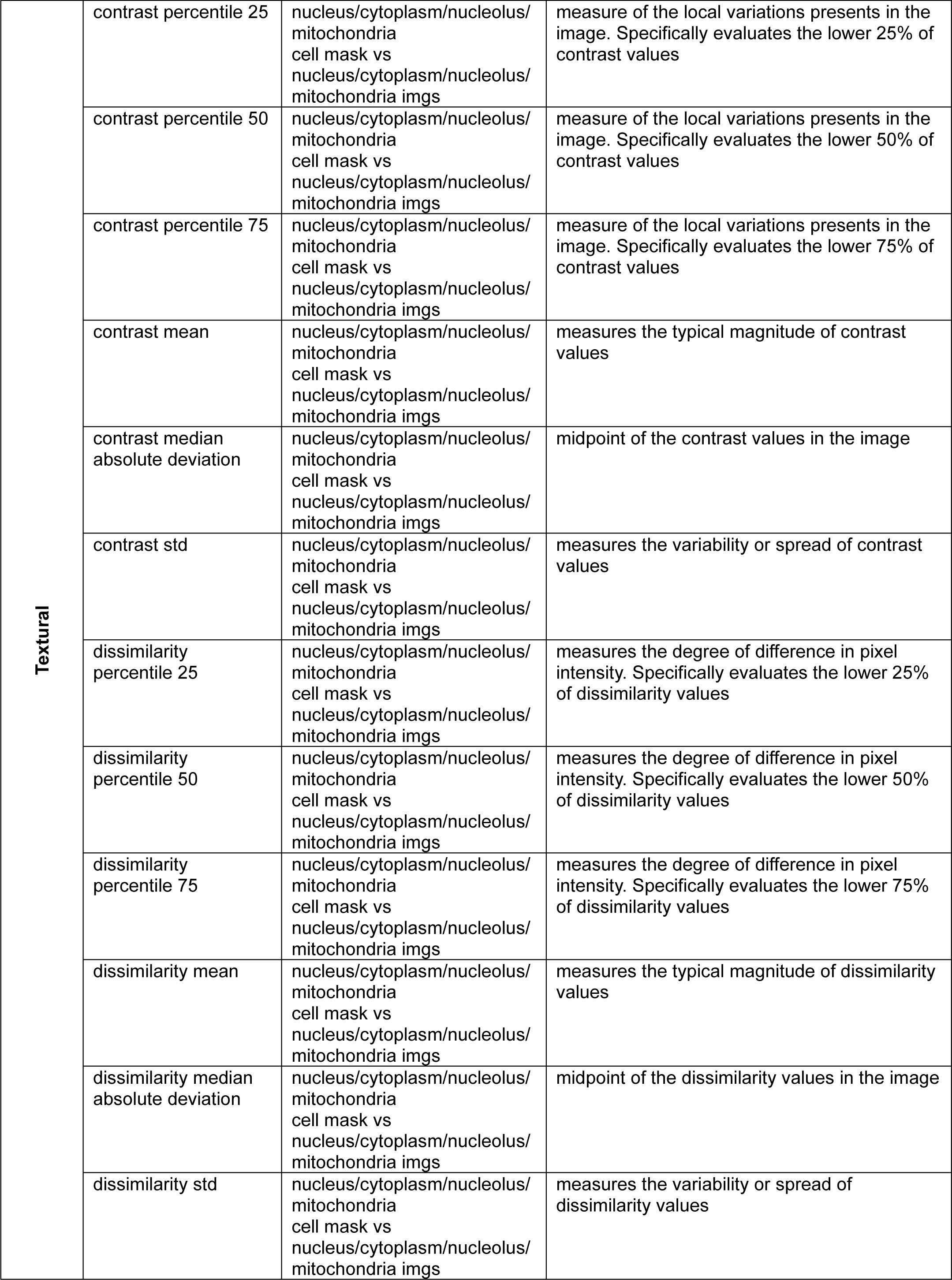
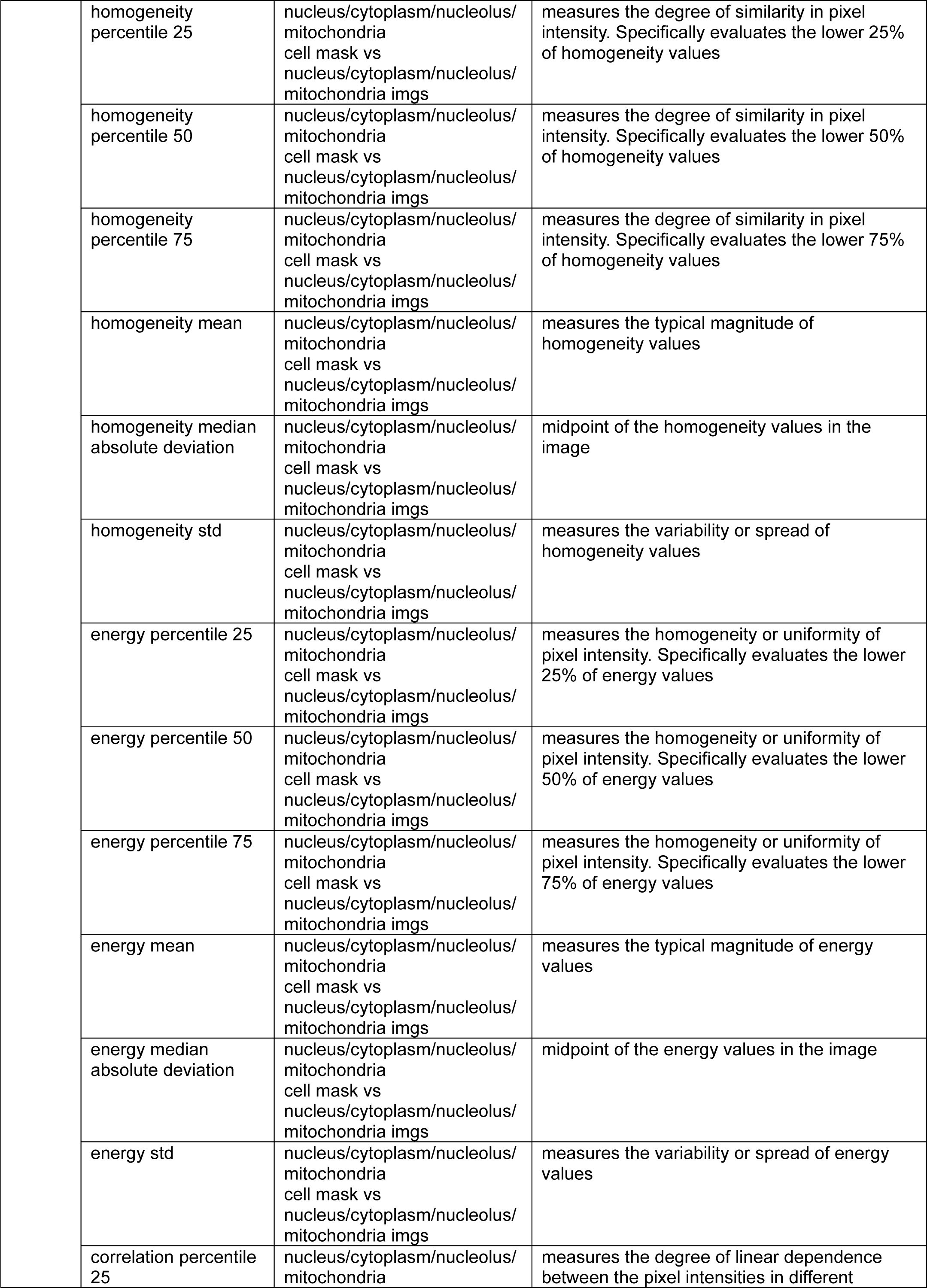
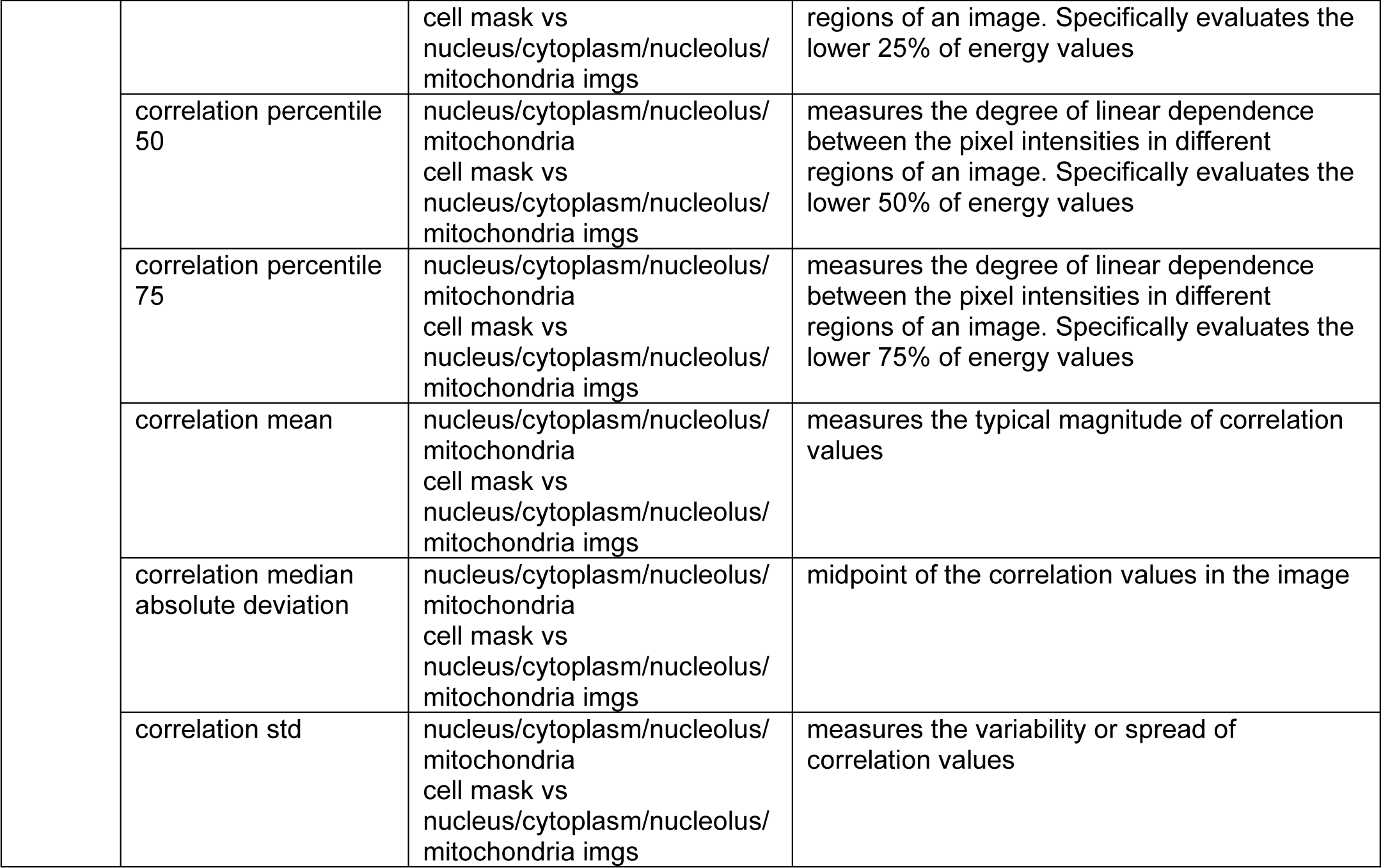
Description of SPACe extracted features.

|  | Feature name | Channel | Description |
| --- | --- | --- | --- |
| Morphology | area | cell/nucleus/cytoplasm/<br>nucleolus/mitochondria | Area of the object (*i.e.* number of pixels of the object scaled by pixel-area) |
|  | convex area | nucleus/cell | Area of the convex hull image, which is the smallest convex polygon that encloses the object |
|  | perimeter | nucleus/cell | Perimeter of object which approximates the contour as a line through the centers of border pixels using a 4-connectivity |
|  | perimeter Crofton | nucleus/cell | Perimeter of object approximated by the Crofton formula in 4 directions |
|  | circularity | nucleus/cell | Circularity defines how closely an object resembles a perfect circle |
|  | efc ratio | nucleus | The EFC ratio measures the regularity of nuclei. A larger ratio indicates more circular or ellipsoidal shapes, while a smaller ratio suggests irregular and complex nuclear shape |
|  | eccentricity | nucleus/cell | Eccentricity of the ellipse having the same second moments as the object. The eccentricity is the ratio of the focal distance (distance between focal points) over the major axis length. The value is in the interval [0, 1). When it is 0, the ellipse becomes a circle |
|  | equivalent diameter area | nucleus/cell | The diameter of a circle with the same area as the object |
|  | Feret diameter max | nucleus/cell | Maximum Feret's diameter computed as the longest distance between points around a region's convex hull contour |
|  | solidity | nucleus/cell | Ratio of pixels in the object to pixels of the convex hull image |
|  | extent | nucleus/cell | Ratio of pixels in the object to pixels in the total bounding box |
| Intensity | Intensity percentile 10 | nucleus/cytoplasm/nucleolus/<br>mitochondria | intensity value below which 10% of the pixel values in the image are found |
|  | intensity percentile 25 | nucleus/cytoplasm/nucleolus/<br>mitochondria | intensity value below which 25% of the pixel values in the image are found |
|  | intensity percentile 75 | nucleus/cytoplasm/nucleolus/<br>mitochondria | intensity value below which 75% of the pixel values in the image are found |
|  | intensity percentile 90 | nucleus/cytoplasm/nucleolus/<br>mitochondria | intensity value below which 90% of the pixel values in the image are found |
|  | intensity median | nucleus/cytoplasm/nucleolus/<br>mitochondria | middle point of the pixel intensity distribution within the image |
|  | intensity mean | nucleus/cytoplasm/nucleolus/<br>mitochondria | average pixel intensity within the image |
|  | intensity median absolute deviation | nucleus/cytoplasm/nucleolus/<br>mitochondria | quantifies the spread or dispersion of pixel intensity values around the median intensity in the image |
|  | intensity std | nucleus/cytoplasm/nucleolus/<br>mitochondria | quantifies the amount of variation or dispersion of pixel intensity values within the image |
| Textural | contrast percentile 25 | nucleus/cytoplasm/nucleolus/<br>mitochondria<br>cell mask vs<br>nucleus/cytoplasm/nucleolus/<br>mitochondria imgs | measure of the local variations presents in the image. Specifically evaluates the lower 25% of contrast values |
|  | contrast percentile 50 | nucleus/cytoplasm/nucleolus/<br>mitochondria<br>cell mask vs<br>nucleus/cytoplasm/nucleolus/<br>mitochondria imgs | measure of the local variations presents in the image. Specifically evaluates the lower 50% of contrast values |
|  | contrast percentile 75 | nucleus/cytoplasm/nucleolus/<br>mitochondria<br>cell mask vs<br>nucleus/cytoplasm/nucleolus/<br>mitochondria imgs | measure of the local variations presents in the image. Specifically evaluates the lower 75% of contrast values |
|  | contrast mean | nucleus/cytoplasm/nucleolus/<br>mitochondria<br>cell mask vs<br>nucleus/cytoplasm/nucleolus/<br>mitochondria imgs | measures the typical magnitude of contrast values |
|  | contrast median<br>absolute deviation | nucleus/cytoplasm/nucleolus/<br>mitochondria<br>cell mask vs<br>nucleus/cytoplasm/nucleolus/<br>mitochondria imgs | midpoint of the contrast values in the image |
|  | contrast std | nucleus/cytoplasm/nucleolus/<br>mitochondria<br>cell mask vs<br>nucleus/cytoplasm/nucleolus/<br>mitochondria imgs | measures the variability or spread of contrast values |
|  | dissimilarity<br>percentile 25 | nucleus/cytoplasm/nucleolus/<br>mitochondria<br>cell mask vs<br>nucleus/cytoplasm/nucleolus/<br>mitochondria imgs | measures the degree of difference in pixel intensity. Specifically evaluates the lower 25% of dissimilarity values |
|  | dissimilarity<br>percentile 50 | nucleus/cytoplasm/nucleolus/<br>mitochondria<br>cell mask vs<br>nucleus/cytoplasm/nucleolus/<br>mitochondria imgs | measures the degree of difference in pixel intensity. Specifically evaluates the lower 50% of dissimilarity values |
|  | dissimilarity<br>percentile 75 | nucleus/cytoplasm/nucleolus/<br>mitochondria<br>cell mask vs<br>nucleus/cytoplasm/nucleolus/<br>mitochondria imgs | measures the degree of difference in pixel intensity. Specifically evaluates the lower 75% of dissimilarity values |
|  | dissimilarity mean | nucleus/cytoplasm/nucleolus/<br>mitochondria<br>cell mask vs<br>nucleus/cytoplasm/nucleolus/<br>mitochondria imgs | measures the typical magnitude of dissimilarity values |
|  | dissimilarity median<br>absolute deviation | nucleus/cytoplasm/nucleolus/<br>mitochondria<br>cell mask vs<br>nucleus/cytoplasm/nucleolus/<br>mitochondria imgs | midpoint of the dissimilarity values in the image |
|  | dissimilarity std | nucleus/cytoplasm/nucleolus/<br>mitochondria<br>cell mask vs<br>nucleus/cytoplasm/nucleolus/<br>mitochondria imgs | measures the variability or spread of dissimilarity values |

|  |  |  |
| --- | --- | --- |
| homogeneity percentile 25 | nucleus/cytoplasm/nucleolus/<br>mitochondria<br>cell mask vs<br>nucleus/cytoplasm/nucleolus/<br>mitochondria imgs | measures the degree of similarity in pixel intensity. Specifically evaluates the lower 25% of homogeneity values |
| homogeneity percentile 50 | nucleus/cytoplasm/nucleolus/<br>mitochondria<br>cell mask vs<br>nucleus/cytoplasm/nucleolus/<br>mitochondria imgs | measures the degree of similarity in pixel intensity. Specifically evaluates the lower 50% of homogeneity values |
| homogeneity percentile 75 | nucleus/cytoplasm/nucleolus/<br>mitochondria<br>cell mask vs<br>nucleus/cytoplasm/nucleolus/<br>mitochondria imgs | measures the degree of similarity in pixel intensity. Specifically evaluates the lower 75% of homogeneity values |
| homogeneity mean | nucleus/cytoplasm/nucleolus/<br>mitochondria<br>cell mask vs<br>nucleus/cytoplasm/nucleolus/<br>mitochondria imgs | measures the typical magnitude of homogeneity values |
| homogeneity median absolute deviation | nucleus/cytoplasm/nucleolus/<br>mitochondria<br>cell mask vs<br>nucleus/cytoplasm/nucleolus/<br>mitochondria imgs | midpoint of the homogeneity values in the image |
| homogeneity std | nucleus/cytoplasm/nucleolus/<br>mitochondria<br>cell mask vs<br>nucleus/cytoplasm/nucleolus/<br>mitochondria imgs | measures the variability or spread of homogeneity values |
| energy percentile 25 | nucleus/cytoplasm/nucleolus/<br>mitochondria<br>cell mask vs<br>nucleus/cytoplasm/nucleolus/<br>mitochondria imgs | measures the homogeneity or uniformity of pixel intensity. Specifically evaluates the lower 25% of energy values |
| energy percentile 50 | nucleus/cytoplasm/nucleolus/<br>mitochondria<br>cell mask vs<br>nucleus/cytoplasm/nucleolus/<br>mitochondria imgs | measures the homogeneity or uniformity of pixel intensity. Specifically evaluates the lower 50% of energy values |
| energy percentile 75 | nucleus/cytoplasm/nucleolus/<br>mitochondria<br>cell mask vs<br>nucleus/cytoplasm/nucleolus/<br>mitochondria imgs | measures the homogeneity or uniformity of pixel intensity. Specifically evaluates the lower 75% of energy values |
| energy mean | nucleus/cytoplasm/nucleolus/<br>mitochondria<br>cell mask vs<br>nucleus/cytoplasm/nucleolus/<br>mitochondria imgs | measures the typical magnitude of energy values |
| energy median absolute deviation | nucleus/cytoplasm/nucleolus/<br>mitochondria<br>cell mask vs<br>nucleus/cytoplasm/nucleolus/<br>mitochondria imgs | midpoint of the energy values in the image |
| energy std | nucleus/cytoplasm/nucleolus/<br>mitochondria<br>cell mask vs<br>nucleus/cytoplasm/nucleolus/<br>mitochondria imgs | measures the variability or spread of energy values |
| correlation percentile 25 | nucleus/cytoplasm/nucleolus/<br>mitochondria | measures the degree of linear dependence between the pixel intensities in different |
|  |  | cell mask vs<br>nucleus/cytoplasm/nucleolus/<br>mitochondria imgs | regions of an image. Specifically evaluates the<br>lower 25% of energy values |
|  | correlation percentile<br>50 | nucleus/cytoplasm/nucleolus/<br>mitochondria<br>cell mask vs<br>nucleus/cytoplasm/nucleolus/<br>mitochondria imgs | measures the degree of linear dependence<br>between the pixel intensities in different<br>regions of an image. Specifically evaluates the<br>lower 50% of energy values |
|  | correlation percentile<br>75 | nucleus/cytoplasm/nucleolus/<br>mitochondria<br>cell mask vs<br>nucleus/cytoplasm/nucleolus/<br>mitochondria imgs | measures the degree of linear dependence<br>between the pixel intensities in different<br>regions of an image. Specifically evaluates the<br>lower 75% of energy values |
|  | correlation mean | nucleus/cytoplasm/nucleolus/<br>mitochondria<br>cell mask vs<br>nucleus/cytoplasm/nucleolus/<br>mitochondria imgs | measures the typical magnitude of correlation<br>values |
|  | correlation median<br>absolute deviation | nucleus/cytoplasm/nucleolus/<br>mitochondria<br>cell mask vs<br>nucleus/cytoplasm/nucleolus/<br>mitochondria imgs | midpoint of the correlation values in the image |
|  | correlation std | nucleus/cytoplasm/nucleolus/<br>mitochondria<br>cell mask vs<br>nucleus/cytoplasm/nucleolus/<br>mitochondria imgs | measures the variability or spread of<br>correlation values |

**Table S2.**
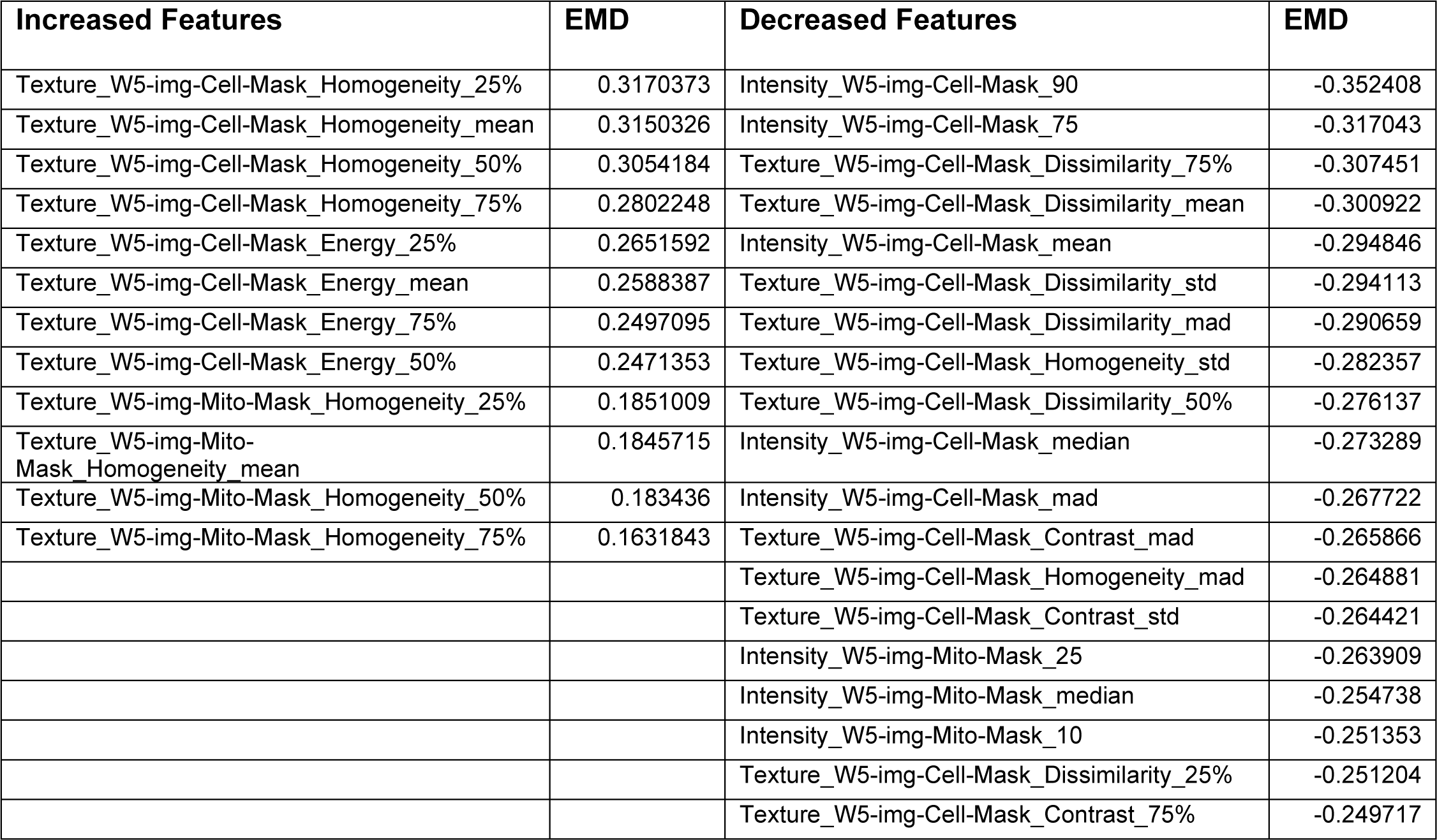
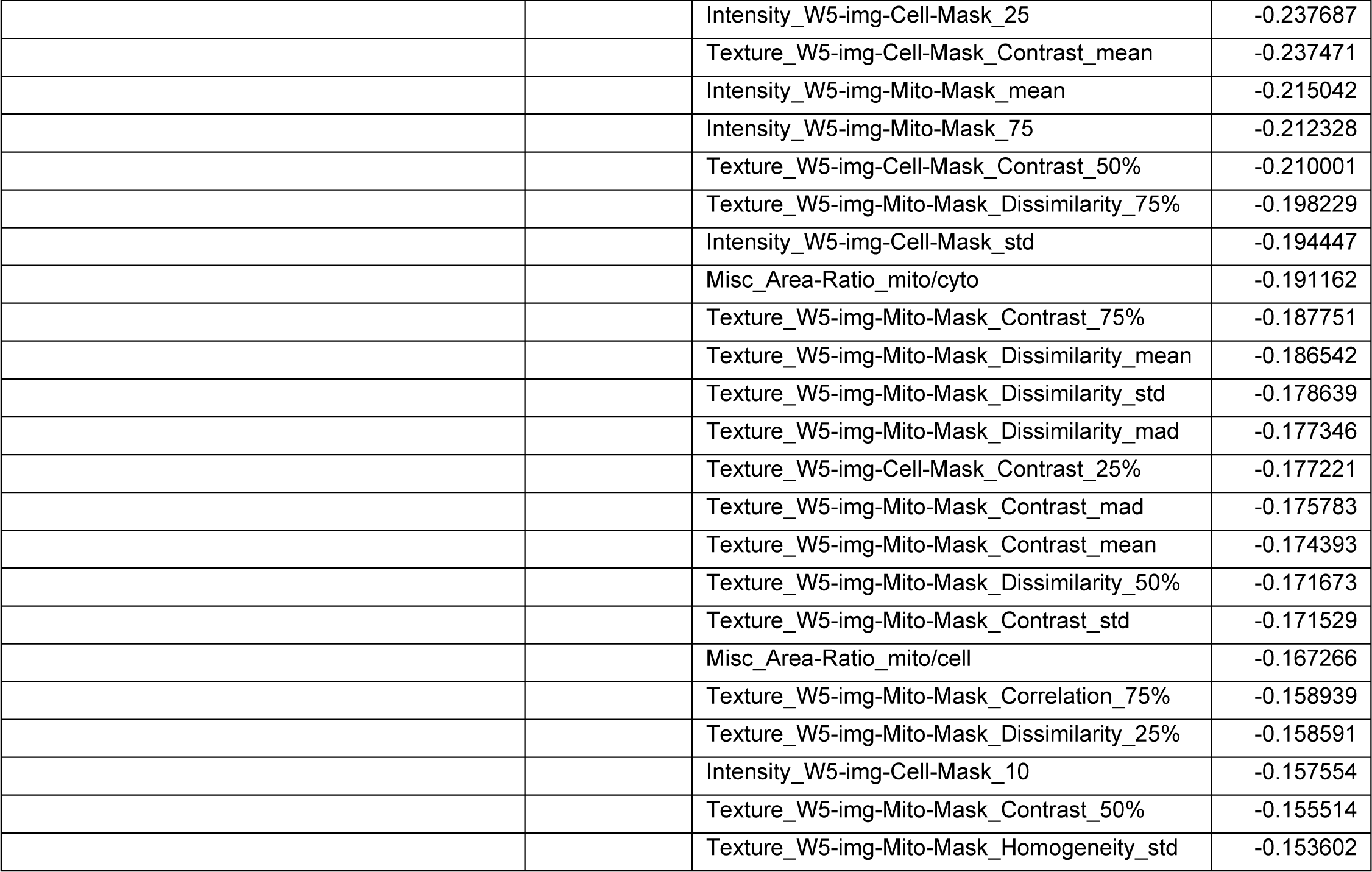
Features included in the berberine chloride consensus fingerprint.

### Supplementary Figure Legends

**Figure S1.**
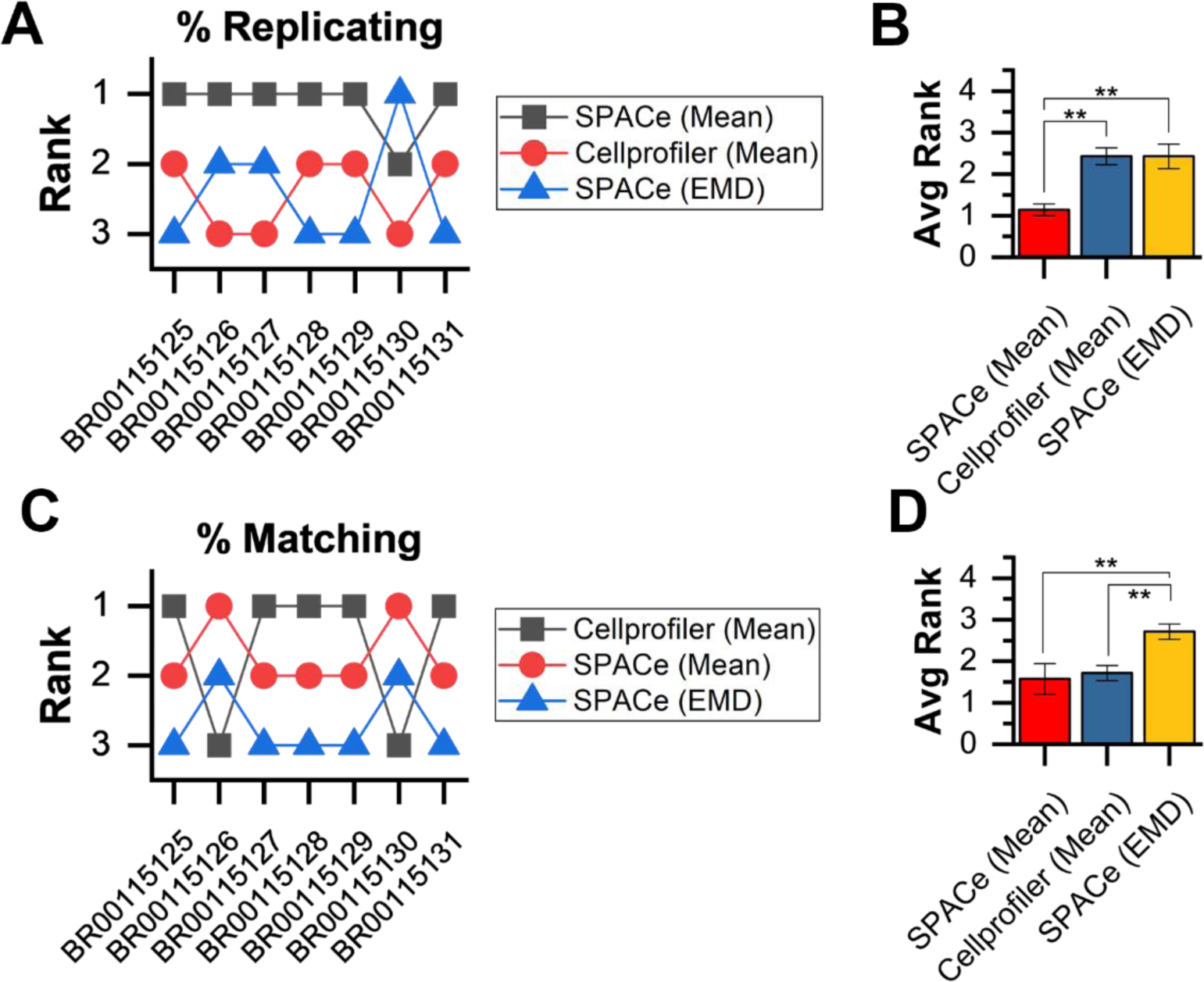
Rank performance of CellProfiler and SPACe analysis of JUMP MOA reference datasets. A) Rank order of percent replicating performance of CellProfiler and SPACe features across JUMP MOA reference datasets. B) Bar chart and statistical analysis of percent replicating performance. C) Rank order of percent matching performance of CellProfiler and SPACe features across JUMP MOA reference datasets. D) Bar chart and statistical analysis of percent matching performance. Brackets and (**) indicates a p-value < 0.01.

**Figure S2.**
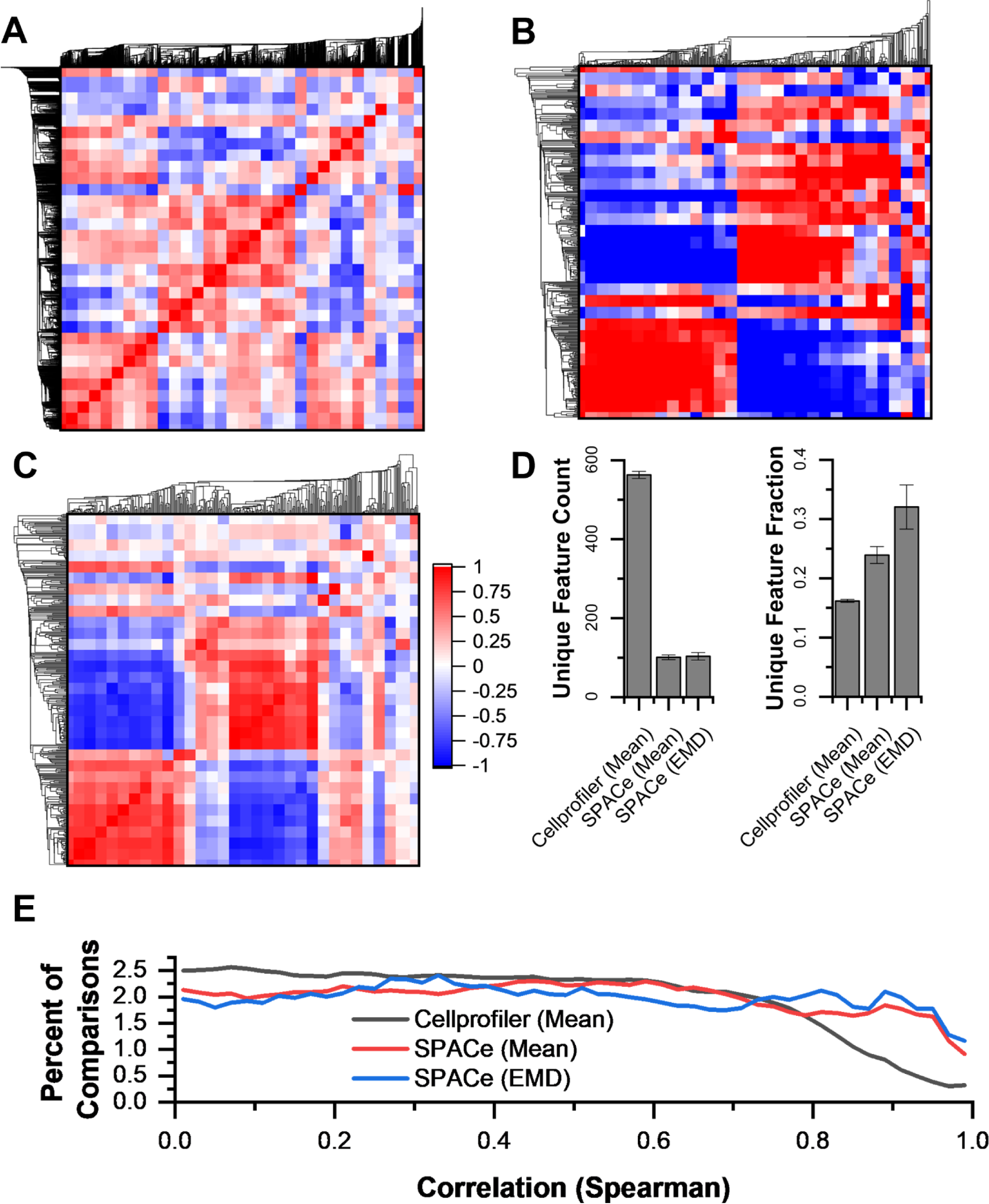
Correlation analysis of extracted features in the JUMP MOA reference datasets. Median correlation (Spearman) matrix across seven JUMP MOA reference datasets for feature extracted using (A) Cellprofiler or SPACe (B, mean; C, EMD). Features are clustered using a Euclidian distance nearest neighbor algorithm. D) Absolute count (left) and fraction (right) of unique features for CellProfiler and SPACe analysis. E) Histogram distribution of correlation values observed between features produced by CellProfiler or SPACe analysis.

**Figure S3.**
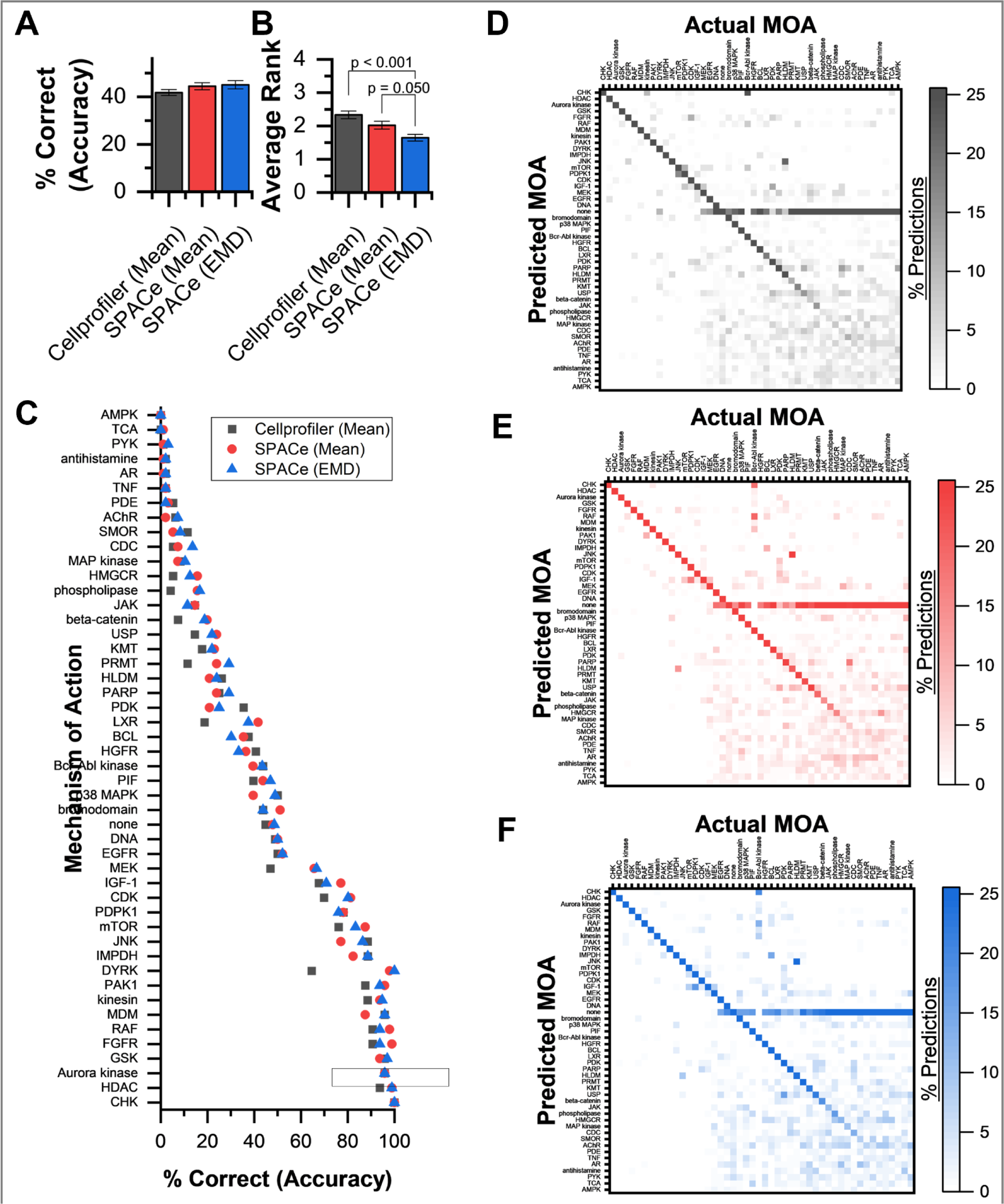
Machine learning model prediction performance using CellProfiler or SPACe extracted features. For six JUMP MOA reference datasets, a set of 5 Random Forest (RF) models were generated to predict MOA using either CellProfiler (Mean), SPACe (Mean), or SPACe (EMD) values. A) Mean and standard deviation of overall accuracy for RF models. B) Mean and standard deviation of model accuracy rank (1 – best, 3 – worst) based on each MOA contained in JUMP reference datasets. Brackets and p-values indicate significant differences observed. C) Average RF model % correct (accuracy) predictions for each MOA contained in the JUMP reference datasets. D-F) Confusion matrices showing the average classification performance of models trained and tested using CellProfiler (Mean), SPACe (Mean), and SPACe (EMD) data, respectively. X and Y axis are ordered based on the order found in (C).

**Figure S4.**
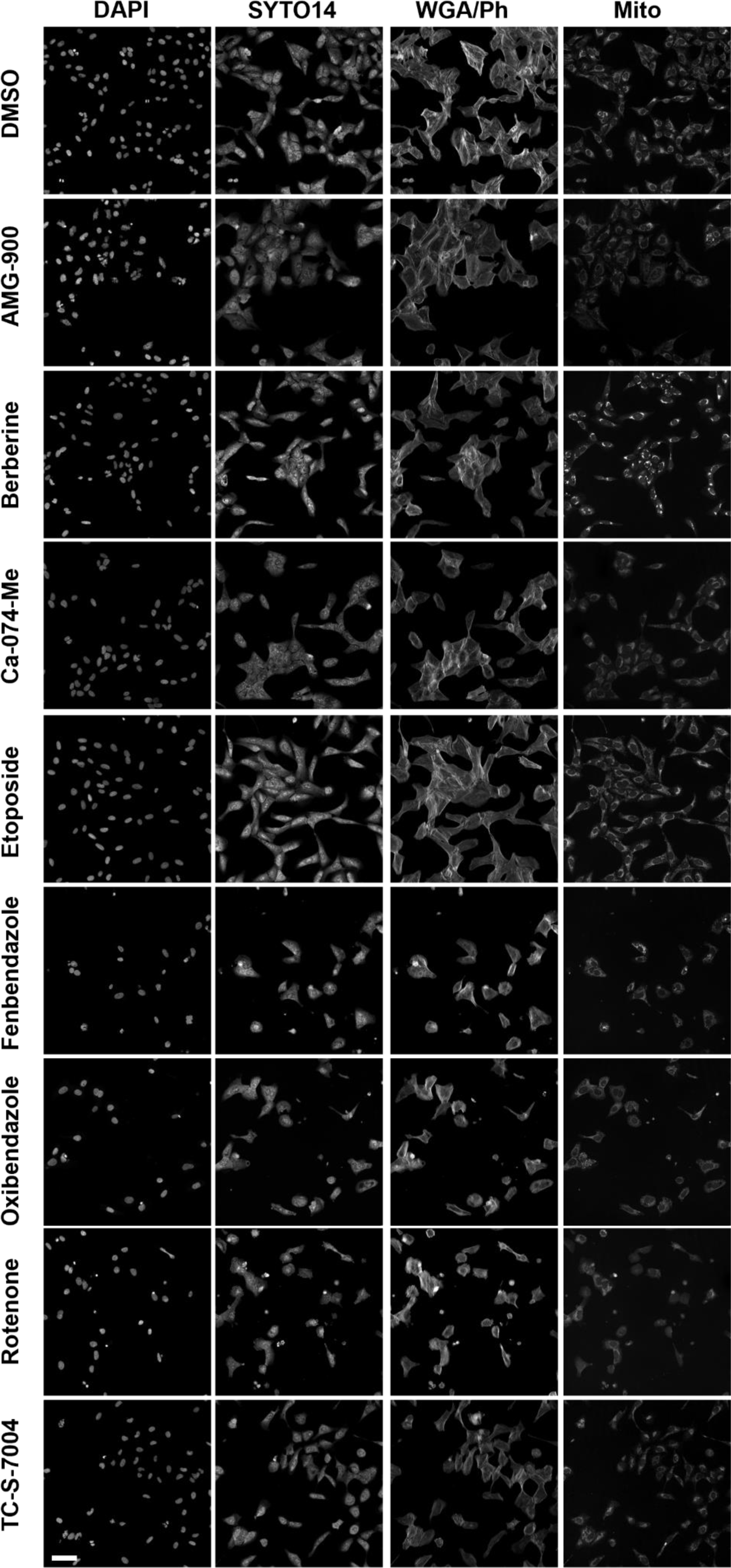
Greyscale images of the treatments described in Figure 2 representing the DAPI, SYTO14, WGA/phalloidin and MitoTracker channels. Scale bar: 50µm.

**Figure S5.**
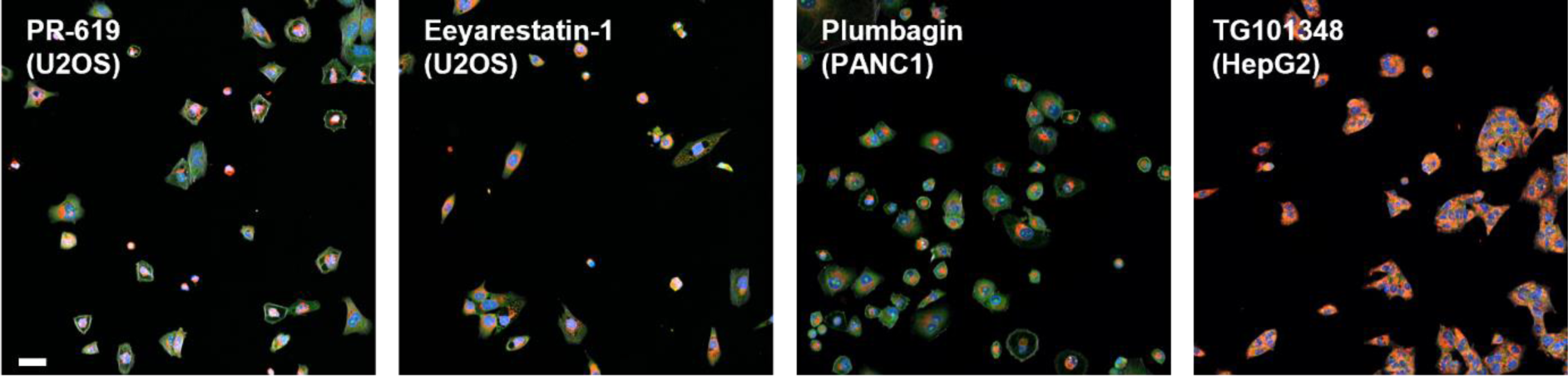
Representative images of the indicated treatments that showed unique phenotypes but also some toxicity in the various cell lines. Scale bar: 20µm.

## Notes

### Competing Interest Statement

The authors have declared no competing interest.

